# Base-pair scale dynamics of a repair helicase on DNA lesions reveal varied damage-sensing mechanisms

**DOI:** 10.64898/2026.03.03.709311

**Authors:** Alice Troitskaia, Tanya Lasitza-Male, Colleen C. Caldwell, Maria Spies, Yann R. Chemla

**Affiliations:** Center for Biophysics and Quantitative Biology, United States; Department of Physics, University of Illinois at Urbana-Champaign, Illinois, United States; Department of Biochemistry, Carver College of Medicine, University of Iowa, Iowa City, Iowa, United States; Department of Physics and Astronomy, and LaserLaB Amsterdam, Vrije Universiteit Amsterdam, The Netherlands

## Abstract

Nucleotide excision repair (NER) is a cellular pathway that removes DNA lesions caused by ultraviolet light and various mutagens. A critical component of the NER machinery is XPD helicase, which unwinds the duplex around the damage, allowing its excision and repair. XPD has also been increasingly implicated in sensing and verifying DNA damage. However, the detailed mechanisms by which XPD responds to DNA damage have to-date remained unclear. Here, we use optical tweezers to perform real-time, high-precision measurements of single molecules of XPD as they encounter a variety of well-defined DNA modifications, including a cyclobutane pyrimidine dimer (CPD), a natural substrate for NER. The observed XPD dynamics reveal different behaviors depending on the damage type and the relative orientations of the DNA fork, damage, and helicase. Most notably, XPD displays an almost complete inability to unwind past a CPD on the translocated strand, and instead exhibits a short pause around the lesion—though not a stall—followed by retreat. Combining the base pair-scale XPD kinetics with structural analyses, we identify two regions of XPD sensitive to DNA modifications.

## Introduction

It is estimated that the genome of each cell experiences up to 10^5^ lesions per day from damage induced by environmental agents or generated endogenously during normal cellular metabolism^1–3^. To maintain genome integrity, cells have evolved a number of mechanisms to sense and repair varying types of DNA damage^4,5^. One such pathway is nucleotide excision repair (NER), which processes a range of lesions: from cyclobutane pyrimidine dimers (CPD) induced by UV irradiation to bulky adducts produced by environmental mutagens^4^. The multi-protein complex TFIIH plays a major role in eukaryotic NER^6–8^, and a key component, the helicase XPD, is vital to NER^9,10^. XPD unwinds the duplex around the DNA lesion, enabling its excision and ultimate replacement with undamaged DNA. Importantly, in addition to unwinding damaged DNA, XPD has been implicated in sensing and verifying DNA damage^11–18^.

XPD belongs to the Superfamily 2B (SF2B) of nucleic acid translocases^19–22^. It comprises a motor core with two RecA-like domains^23–25^, HD1 and HD2, that translocates in a 5’ → 3’ direction along DNA^11,26–28^ (**Fig. 1a**). XPD also has two insertions in HD1^23–25:^ an Arch domain and an FeS domain that contains a 4Fe-4S cluster essential for its helicase activity^19^. During unwinding, the duplex is separated near the pore formed by the Arch, FeS, and HD1 domains^29^. The translocated strand of the unwound DNA fork passes through this pore, through a binding cleft spanning HD1 and HD2, exiting out the back of HD2 where it makes additional contacts^27–32^. Extended DNA interactions within XPD have pointed to a structural basis for a damage-sensing role. In particular, a lesion sensor pocket was proposed near the entrance to the pore^15^, supported by biochemical^15,27,28^ and biophysical^18^ data, with further evidence from structural^29,31^ and computational^33,34^ studies showing contacts between the translocated strand and residues in this sensor pocket.

**Figure 1.**
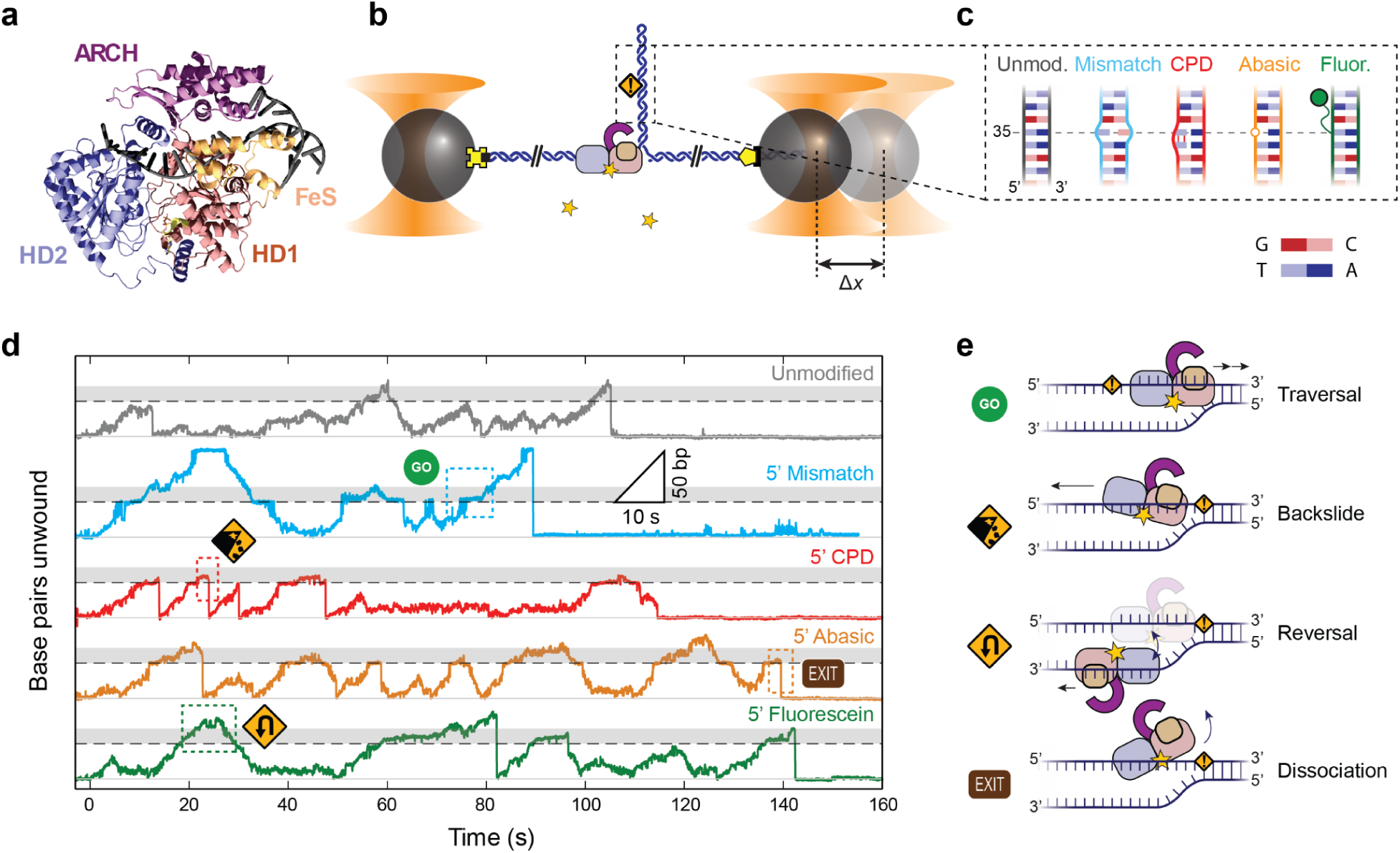
XPD responds differentially to DNA modifications. (**a**) Structure of XPD helicase (PDB 8rev), showing its two motor domains (HD1, salmon; and HD2, blue), accessory Arch (purple) and FeS domains (orange), and its interactions with the translocated strand of a DNA fork. (**b**) Schematic of the hairpin unwinding assay. A single molecule of DNA containing an 86-bp hairpin (dark blue) is tethered between two micron-sized beads held in optical traps (orange cones) via biotin-streptavidin (yellow cross) and digoxigenin-antibody (yellow pentagon) attachments. A single XPD helicase binds to a 10-nt ssDNA site at the 5’ end of the hairpin and unwinds it upon addition of ATP (yellow star). Unwinding of each hairpin base pair increases the end-to-end extension of the tether at a constant force. A DNA modification (orange warning sign) resides 35 bp from the base of the hairpin. (**c**) Schematics of the 5’ set of DNA constructs, showing the unmodified ‘core’ sequence with uniform base pairing stability (gray) and those containing a modification on the 5’ strand: single-base mismatch (light blue), 19yclobutene pyrimidine dimer (CPD; red), abasic site (orange), and fluorescein (green). (**d**) Representative unwinding traces on each DNA construct (same colors as c), showing stereotypical behaviors upon encountering a DNA modification. Traces are offset vertically for clarity. For each trace, the light gray line indicates the position of the base of the hairpin and the dashed line the position of the modification; shaded areas indicate XPD’s interaction region with the modification. € Schematic of XPD behaviors during encounters with modifications: traversal, backslide, reversal, and dissociation.

Biochemical evidence for XPD DNA damage detection has been accumulating for years^11–18,35^, although not without controversy. While various studies have found that DNA modifications inhibit XPD homologs from the archaea *Ferroplasma acidarmanus* (FacXPD)^14,15^ and *Thermoplasma acidophilum* (taXPD)^16^ and from eukaryotes *C. thermophilum* (ctXPD)^17^ and yeast (Rad3)^12,13,36^, another study reported that the archaeon *Sulfolobus acidocaldarius* (SacXPD) could unwind damage-containing duplexes to the same extent as undamaged duplexes^37^. Differences between homologs and experimental conditions could account for these seemingly contradictory results^14,16,38^. In systems where DNA damage was observed to affect XPD activity, there remains a lack of consensus on the strand specificity of damage detection and how it may differ by type of damage^12–17,35^. There also remain disagreements over the nature of the inhibition by DNA damage. Biochemical studies of FacXPD^14,15^ and AFM imaging studies of taXPD^16^ and ctXPD^17^ point to the formation of long-lived complexes on damaged DNA. However, single-molecule fluorescence studies of FacXPD^18^ showed that its average residence time on a DNA bubble containing a CPD site was comparable to that on undamaged DNA, arguing against stable complex formation.

The mechanisms by which XPD may sense damage remain unclear, highlighting a need for real-time, high precision measurements of XPD dynamics on damage-containing DNA. Here, we use optical traps to measure the unwinding activity of XPD on DNA with a well-defined damage site, using FacXPD, an established model for biochemical^14,15,39,40^ and biophysical studies^18,41–43^ of XPD, with a damage sensing mechanism thought to resemble that of human XPD^15^. The high spatiotemporal resolution of the measurements allows us to detect base pair-scale dynamics of individual molecules of XPD as they encounter a variety of DNA modifications, including a CPD (an important substrate for NER^4^), fluorescein (a proxy for bulky extrahelical adducts^37,16,17^), and an abasic site (the outcome of frequent endogenous depurination^3^). Our measurements show that XPD senses DNA damage, with a sensitivity that depends on the damage type and the relative orientations of the helicase, damage, and DNA fork. Sensing is manifested by increased residence times around the lesion and, in some cases, by an impaired ability to bypass the lesion site. The base-pair scale resolution of our measurements allows us to identify two regions of the protein that interact with damage, thus providing new insights into the structural mechanisms of damage sensing.

## Results

### XPD does not unwind past a CPD on the translocated strand but bypasses other modifications

We measured the duplex unwinding activity of individual XPD molecules on DNA containing different modifications using a previously described optical trap assay^41^. Briefly, dual-trap optical tweezers^44–46^ were used to hold two polystyrene beads between which a single DNA molecule was tethered (**Fig. 1b**). The DNA comprised an 86-bp hairpin containing the modification of interest and two flanking ∼1.5-kb double-stranded (ds)DNA spacers. A 10-dT single-stranded (ss)DNA segment on the 5’ end of the hairpin served as a loading site for the helicase (see **Methods**). Based on structural data, XPD’s footprint on ssDNA is around 11-14 nt^29,31^ (see also **Supplementary Methods**), although we previously showed that 10 nt is sufficient for binding a single XPD^41,42^.

We synthesized a set of DNA substrates containing different types of modifications designed to test XPD’s ability to negotiate disruptions to the duplex and to probe damage-specific interactions. Previously we found that XPD unwinding was strongly affected by the local base pairing stability^47–49^ of the DNA sequence^41^ (see **Methods**). We therefore synthesized ‘core’ DNA substrates with uniform base pairing stability to minimize sequence-dependent effects (**Fig. 1c**, gray schematic; **Supplementary Fig. 1**, see **Methods**). Into these core substrates we inserted the following modifications, some the subjects of prior studies on XPD damage sensing^13–18,37^: (i) a single-base pair mismatch, which serves as a control substrate for measuring XPD activity on DNA with a disrupted base pair but unmodified base (**Fig. 1c**, light blue schematic); (ii) a CPD (red); (iii) an abasic site, a form of damage that may interfere with XPD-DNA interactions^13^ (orange); and (iv) a fluorescein dye attached via a 6-carbon linker (green). Two sets of substrates were synthesized with the modification on either strand of the hairpin stem (labeled the 5’ and 3’ set, respectively; **Supplementary Fig. 2a, d**, see **Methods**), to compare XPD’s response to damage on the translocated or displaced strand^13–17^. In each case, the modification was incorporated at a single site on the hairpin stem (base pair 35 as measured from the stem base; see **Methods**).

We verified the proper construction of each substrate by measuring its elastic behavior with a force-extension curve (FEC), as shown in **Supplementary Figure 2b, e** (see **Methods**). In contrast to the uniform core sequences (gray curves), the FECs of hairpins containing modifications display a rip in the middle of the unfolding transition (light and dark blue, red, orange, and green curves), due to local disruption in duplex stability around the modifications^50–52^. Analyzing the unfolding transitions, we ranked the degree of disruption from each modification (**Supplementary Fig. 2c, f**). The CPD destabilizes the duplex less than the mismatch or abasic site, consistent with findings that its perturbation of local DNA structure is subtle^50^; fluorescein destabilizes the duplex the least.

All XPD unwinding experiments were carried out in a laminar flow chamber^44,45^ (**Supplementary Fig. 3**) that maintained two parallel streams, one containing a saturating concentration of ATP^38,40^, the other XPD with ATP-γS (see **Methods**). In a typical measurement, we tethered a DNA molecule between the trapped beads in the ATP stream, moved into the stream containing XPD for ∼1 min, allowing a single molecule of XPD to bind, and returned to the ATP stream to initiate unwinding of the hairpin. The absence of helicase within the ATP stream ensured that all detected activity resulted from the single molecule of XPD pre-loaded to the DNA hairpin. We monitored the DNA fork position with near-base pair resolution from the change in extension of the tether, as each unwound base pair (bp) released 2 nucleotides (nt) (see **Methods**). Active force feedback maintained a constant tension across the DNA of ∼12 pN, below the mechanical unfolding force of the hairpin (**Supplementary Fig. 2**).

Figure 1d shows representative time traces of XPD unwinding hairpins from the 5’ set of substrates. In line with previous measurements^41,42^, a molecule of XPD could unwind DNA in multiple ‘bursts’, repeated cycles of unwinding followed by fork regression toward the hairpin base. We focused our analysis on XPD bursts at the modification site (Fig. 1d, site at 35 bp indicated by dashed lines) and a distance beyond corresponding to XPD’s footprint (35-49 bp, indicated by shaded areas), presumably where XPD was interacting with the modification. We observed various behaviors as XPD encountered the modification (Fig. 1e schematics): (i) traversals (e.g. Fig. 1d, light blue dashed box), during which XPD advanced past the modification by >14 nt, successfully clearing it; (ii) backslides (red dashed box), which we believe represent XPD partially disengaging from the translocated strand and retreating down the hairpin stem^41^; (iii) reversals (green dashed box), which we interpret as XPD switching to the opposing strand to translocate down the stem, allowing the hairpin to rezip gradually behind it^42^; or (iv) dissociations, marked by total cessation of unwinding (orange dashed box).

Figure 2a shows XPD’s traversal probability vs position on the hairpin stem for every modification (**Methods**). The traversal probability on the unmodified DNA (gray) drops with position due to intrinsic limits to XPD’s processivity. The survival profile is similar for the mismatch (light blue) and, remarkably, for the bulky fluorescein addition (green), indicating that XPD can bypass these modifications with the same probability as unmodified duplex. In contrast, the traversal probability on the abasic site (orange) drops off faster, indicating that the damage impedes XPD unwinding. The effect of the CPD (red) is the most dramatic; practically no bursts reached further than 45 bp. Figure 2b displays the probability of each type of XPD behavior upon encountering the modification. XPD encounters with the CPD almost always ended with a backslide or with dissociation, while reversals were rarely observed.

**Figure 2.**
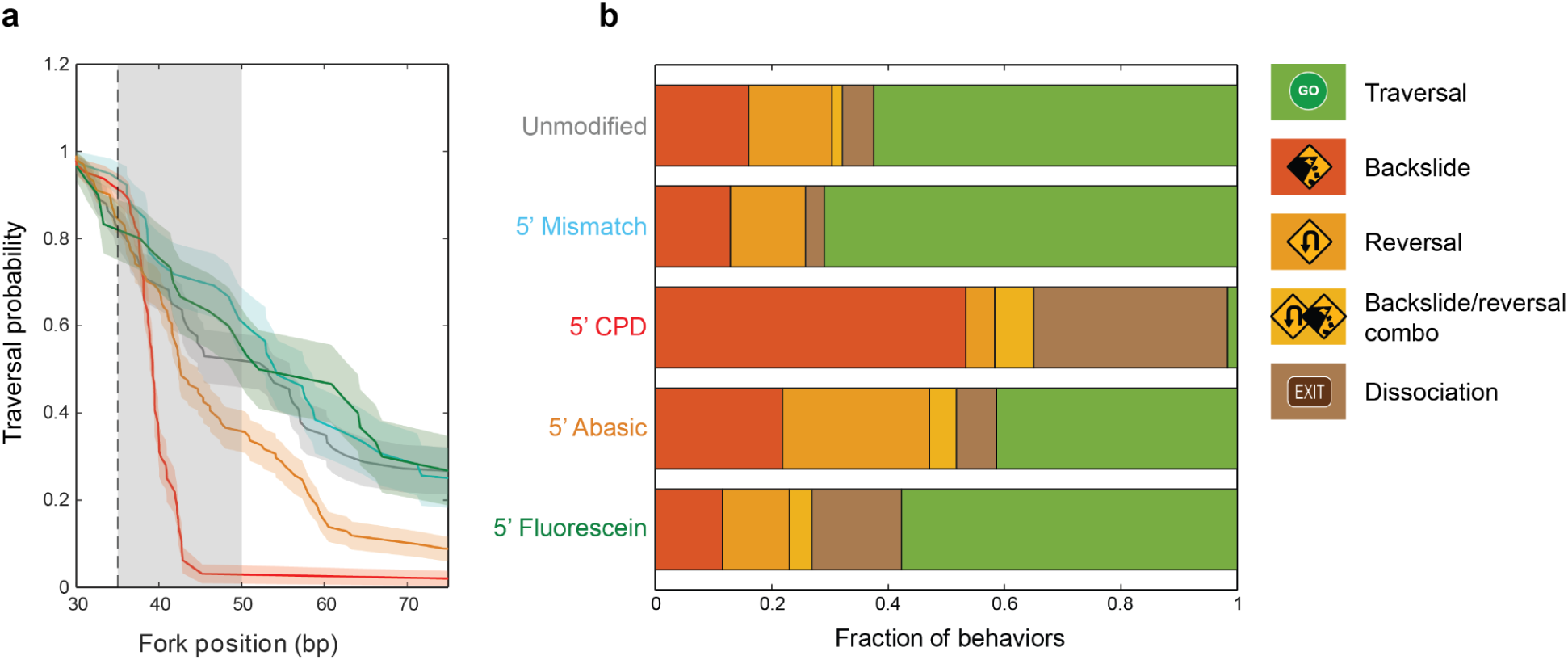
XPD behaviors upon encounters with 5’-strand modifications during unwinding. (**a**) Traversal probabilities past the modification for each DNA construct (same colors as Figure 1). Shaded regions denote s.e.m. The modification site and interaction region are denoted by the dashed line and gray shaded area. (**b**) Fraction of observed behaviors as XPD encounters each 5’ modification: traversal (green), backslide (dark orange), reversal (orange), backslide/reversal combination (yellow), and dissociation (gray). Only bursts that reached at least 30 bp were selected for this analysis.

We repeated the above measurements with hairpins containing the DNA modifications on the opposite, displaced strand of the stem (3’ set; **Supplementary** Fig. 4a). As shown in representative traces on this set of substrates, XPD was able to unwind past the modification site at 35 bp in all cases—including, notably, the CPD (**Supplementary** Figure 4b). Moreover, the traversal probabilities for the unmodified and modified substrates were all nearly indistinguishable (**Supplementary** Fig. 4c), indicating that XPD did not sense modifications on the displaced strand.

### XPD exhibits two different pauses upon unwinding encounters with DNA modifications

Close inspection of the representative traces in Figure 1d reveals pauses around the modifications. For all substrates containing translocated-strand modifications (5’ set) and the unmodified core sequence, we determined the average residence time spent at each position on the hairpin stem (**Supplementary** Fig. 5) to analyze the kinetics of XPD’s encounters with the modifications. The dynamics of modified DNA substrates themselves must be considered to interpret the kinetics correctly. As our assay measures the position of the DNA fork, not necessarily of XPD itself, we cannot distinguish between unwinding by XPD and spontaneous hairpin unfolding. In Figure 1d, the time traces on modified DNA display a characteristic feature in which the duplex rapidly unfolded up to the modification site at 35 bp (e.g. light blue trace at *t* ≈ 51 s). As shown in the example unwinding traces in Figure 3a, when XPD approached the destabilizing modification (Fig. 3b, upper schematic; step 1), the intervening stretch of duplex became sufficiently short as to be unstable, leading to rapid unfolding up to the modification site (step 2). Unfolding was followed by a pause while XPD caught up to the fork and encountered the modification (step 3). Figure 3c displays the residence times on modified hairpins measured relative to those on the core sequence. This sequence of steps is manifested in the decrease in relative residence times at 20-30 bp, followed by the peak near 35 bp (and 43 bp for fluorescein, as explained below; Fig. 3c, dark arrows).

**Figure 3.**
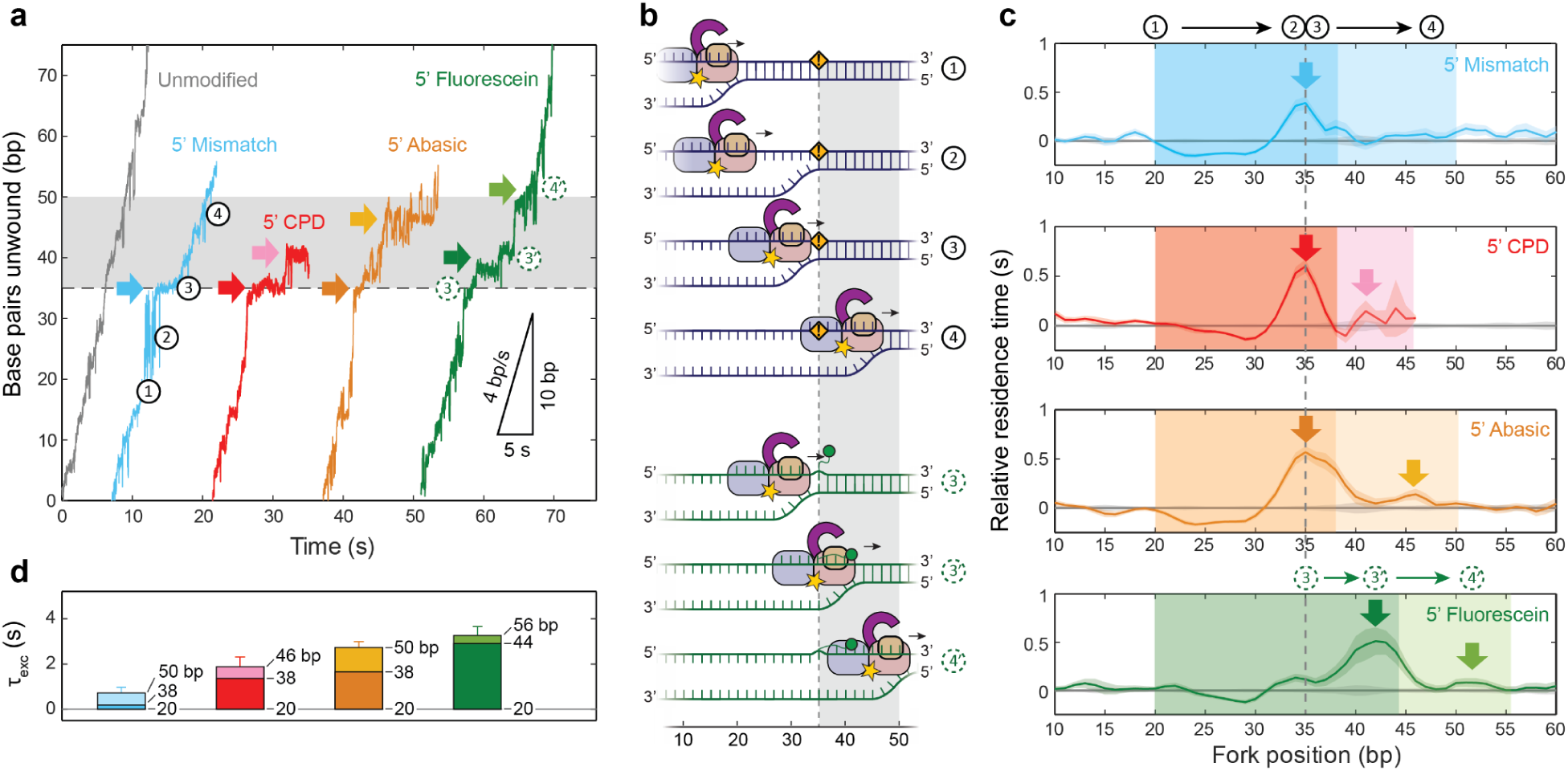
XPD unwinding kinetics on 5’-strand DNA modifications. (**a**) Example traces of XPD unwinding DNA containing 5’-strand modifications (same colors as in Figure 1). Sequence of events: (1) XPD approaches the modification, (2) the hairpin spontaneously unfolds up to the modification due to its destabilizing effect, (3) XPD catches up to the fork leading to an observed pause (dark arrows), and (4) eventually passes the modification. For the fluorescein substrate (green) the sequence of events is altered: (3) XPD approaches the fluorescein modification and (3’) stretches the fluorescein linker, leading to a shift in pause location (dark green arrow). Light arrows indicate second pauses observed within XPD’s interaction region with the modifications. (**b**) Schematics for events (1)-(4) during an XPD encounter with a DNA modification. (**c**) Average residence time vs DNA fork position on the mismatch, CPD, abasic, and fluorescein substrates (top to bottom), measured relative to that on unmodified DNA (gray). Shaded areas around the average times denote standard error of the mean (s.e.m.). Arrows indicate pauses as in a. (**d**) Excess XPD traversal times, *τ_exc_*, for different substrates. *τ_exc_*determined by integrating the relative residence times over the destabilization region (dark shaded rectangles in c) and over the secondary interaction region of XPD with the modification (light shaded rectangles in c); the integration ranges in bp are indicated by each bar. Error bars are determined from s.e.m. of residence times of positions included in integration.

The pause at the modification site represents two kinetic events: XPD catching up to the fork and pausing due to damage-specific interactions. To quantify the damage-specific contribution to pausing, we integrated the relative residence times over the destabilized region through the modification site (20-38 bp). Figure 3d displays this ‘excess’ traversal time, τ*_exc_*, through the modification for each substrate (darkly shaded bars). For the mismatch, τ*_exc_*was nearly zero. The overall decrease in residence time due to spontaneous hairpin opening is compensated by the increase from the pause at the modification, expected if XPD translocated at the same rate as it unwound unmodified DNA and did not slow down at the modification site. (Note that the mismatch substrate was designed so that the sequence of the translocated strand was identical to that of the unmodified substrate; see **Methods**.) In contrast, XPD exhibited excess traversal times of τ*_exc_*= 1.4 ± 0.2 s and 1.7 ± 0.2 s on the CPD and abasic substrates, respectively. These durations indicate that a CPD or abasic site jams XPD upon encounter. For fluorescein, the peak in residence time occurred at 42 bp, offset +7 bp from the modification site (Fig. 3c, dark green arrow). We attribute this offset to the linker through which fluorescein is attached to the base. Like the CPD and abasic sites, fluorescein jammed XPD, resulting in a pause, but only when the linker was fully stretched (Fig. 3b, green schematic, step 3’). We estimated the excess traversal time by integrating relative residence times through the offset peak (20-44 bp), giving τ*_exc_*= 2.9 ± 0.4 s. These excess times correspond to damage-specific pausing as XPD’s front end first encountered the damage (Fig. 3b, step 3 or 3’).

XPD exhibited additional pausing as it unwound further, as the modification made its way through the helicase interior. As shown in Figure 3a and c, a second, ‘internal’ pause at 46 bp is observed for the abasic DNA and at ∼41 bp for the CPD (light arrows). Calculating the excess traversal time over the region encompassing the lesion’s passage through XPD (38-50 bp, at which point the back end of XPD would last make contact with the modification) gives τ*_exc_* = 1.0 ± 0.3 s for the abasic site compared to 0.5 ± 0.3 s for the CPD. For fluorescein, we observe a pause at ∼53 bp, which contributes an excess time of τ*_exc_*= 0.4 ± 0.3 s (integrated over the offset region 44-56 bp). As above, this pause position likely corresponds to that at 46 bp for abasic DNA, shifted +7 bp due to stretching of the linker. The existence of a second pause suggests that important interior contacts, ∼11 nt from XPD’s front end, are altered when XPD encounters modified DNA. **Tables 1, 2** summarize the pause durations and positions, respectively.

**Table 1.** Excess traversal times during XPD encounters with DNA modifications. *Off-strand, ^a^For 5’ CPD, ranges are 20-38 bp and 38-46 bp; for 5’ fluorescein, ranges are 20-44 bp and 44-56 bp. ^b^For 3’ fluorescein, ranges are 45-35 bp and 35-20 bp.

|  | 5' Unwinding |  | 3' Reziping |  |
| --- | --- | --- | --- | --- |
| Pause<br>Mod. | Frontal (s)<br>(20-38)<sup>a</sup> | Internal (s)<br>(38-50 bp)<sup>a</sup> | Frontal (s)<br>(50-42 bp)<sup>b</sup> | Internal (s)<br>(42-20 bp)<sup>b</sup> |
| Mismatch* | 0.2 ± 0.2 | 0.5 ± 0.3 | 0.1 ± 0.2 | 0.4 ± 0.2 |
| CPD | 1.4 ± 0.2 | 0.5 ± 0.3 | 0.5 ± 0.2 | 2.9 ± 0.4 |
| Abasic | 1.7 ± 0.2 | 1.0 ± 0.3 | -0.2 ± 0.3 | 0.7 ± 0.3 |
| Fluorescein | 2.9 ± 0.4 | 0.4 ± 0.3 | 1.1 ± 0.3 | 0.5 ± 0.2 |

**Table 2.** Measured and predicted pause positions. XPD footprint *f_X_* = 14 nt; fluorescein linker length *L_f_* = 7; internal damage sensor packet location, relative to XPD’s front end, Δ = 11 nt.

|  |  | 5' Unwinding |  | 3' Reziping |  |
| --- | --- | --- | --- | --- | --- |
| Pause<br>Mod. |  | Frontal (bp) | Internal (bp) | Frontal (bp) | Internal (bp) |
| CPD | Pred. | 35 | 35 + Δ = 46 | 35 + f<sub>X</sub> = 49 | 35 - Δ + f<sub>X</sub> = 38 |
|  | Meas. | 35 | -- | 47 | 37 |
| Abasic | Pred. | 35 | 35 + Δ = 46 | 35 + f<sub>X</sub> = 49 | 35 - Δ + f<sub>X</sub> = 38 |
|  | Meas. | 35 | 46 | -- | 37 |
| Fluor. | Pred. | 35 + L<sub>f</sub> = 42 | 35 + Δ + L<sub>f</sub> = 53 | 35 + f<sub>X</sub> - L<sub>f</sub> = 42 | 35 - Δ + f<sub>X</sub> - L<sub>f</sub> = 31 |
|  | Meas. | 42 | 52 | 42 | 31 |

We repeated this analysis for XPD unwinding with DNA modifications on the displaced strand (3’ set). As shown in **Supplementary** Fig. 6a, relative residence times showed a similar decrease around 20-30 bp as the fork approached the modification, due to its destabilizing effects. However, the peaks at the modification site, 35 bp, were lower than that for the translocated strand modifications, indicating that XPD was not slowed down when encountering modifications on the displaced strand. Indeed, there was no appreciable pausing over the 20-50 bp interaction region for any modification (**Supplementary** Fig. 6b).

### XPD’s response to DNA modifications differs with a reversed fork-XPD orientation

As discussed above and as reported previously^42^, we routinely observed reversals during bursts (e.g. Fig. 1d gray trace, *t* = 60-65 s), where XPD switches to the opposing strand of the hairpin stem and translocates away from the DNA fork, allowing it to rezip gradually (Fig. 1e). In those cases where XPD unwound past the modification site and then reversed direction, we could study XPD-damage interactions with a reversed fork-XPD geometry.

We first analyzed XPD’s behavior on the 3’ set of substrates, for which the DNA modification resided on the strand translocated during rezipping (e.g. **Supplementary** Fig. 4b, green trace, *t* = 20-30 s). As shown in the plot of traversal probability vs. position in **Supplementary** Fig. 7, XPD could bypass the abasic site, mismatch, and fluorescein with similar ease during rezipping as it did during unwinding. Remarkably, however, XPD could also overcome the CPD site in this configuration. Thus, the response to this type of damage is orientation-dependent, and the CPD does not impose an absolute block. A key difference between the two orientations is that the DNA fork forms at the back end of XPD during rezipping, with fork regression in the same direction as XPD’s translocation down the hairpin stem (see **Discussion**).

In Figure 4a, example traces of hairpin rezipping show that modifications induced pausing, similarly to unwinding. The mean residence times vs fork position measured relative to those on unmodified DNA are shown in Fig. 4c for modified DNA. (The mismatch substrate for these measurements was altered so that the sequence of the translocated strand during XPD rezipping was identical to that of the unmodified substrate; see **Methods**.) Since the geometry is reversed during rezipping, the DNA fork, whose position is monitored in our assay, resides a distance corresponding to XPD’s footprint away from its front end (Fig. 4b, upper schematic, step 1). For the CPD, a pause is observed at ∼47 bp (Fig. 4c, dark arrow), consistent with XPD’s front end sensing the damage at 35 bp given a footprint of ∼11-14 nt^29,31^. On the other hand, XPD showed no pausing on abasic DNA at this position. For fluorescein, a pause was seen at ∼42 bp (Fig. 4c, dark arrow). We again attribute this shift to stretching of the fluorescein linker, eventually jamming the front end of XPD. We integrated the relative residence times over this region to determine the excess traversal time. For the CPD, τ*_exc_* = 0.5 ± 0.2 s (integrating over the region 50-42 bp); for fluorescein, τ*_exc_* = 1.1 ± 0.3 s (45-35 bp, Fig. 4c, darkly shaded areas).

**Figure 4.**
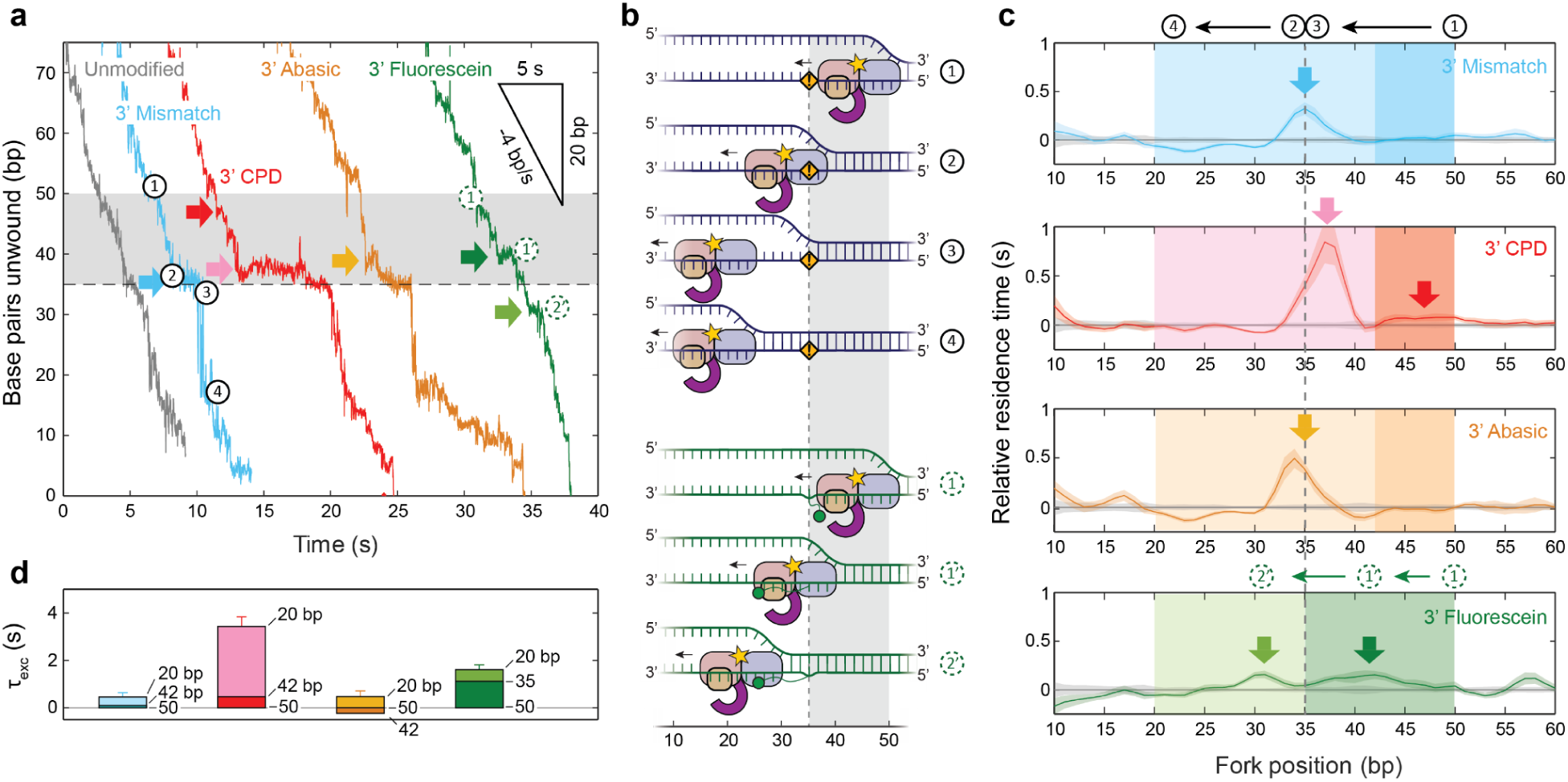
XPD rezipping kinetics on 3’-strand DNA modifications. (**a**) Example traces of XPD rezipping DNA containing 3’-strand modifications (same colors as in Figure 1). Sequence of events: (1) XPD reaches the modification, (2) translocates past the modification, (3) the hairpin remains unfolded up to the modification due to its destabilizing effect, leading to an observed pause, and (4) after XPD vacates enough nucleotides, the DNA fork regresses down to the back end of XPD. For the fluorescein substrate, the schematics (green), the sequence of events is altered: (1) XPD approaches the fluorescein modification and (1’) stretches the fluorescein linker, leading to a shift in pause location. (**b**) Schematics for events (1)-(4) during an XPD encounter with a DNA modification. (**c**) Average residence time vs DNA fork position on the mismatch, CPD, abasic, and fluorescein substrates, measured relative to that on unmodified DNA during XPD rezipping activity. Shaded areas denote standard deviation. (**d**) Excess XPD traversal times, *τ_exc_*, for different substrates over the initial (dark shaded bar) and secondary (lightly shaded) interaction regions with the modification; the integration ranges in bp are indicated by each bar. Error bars are determined from s.e.m..

XPD exhibited additional pausing once past its first encounter with the modification. On mismatched DNA, we observed a pause at the modification site position, 35 bp, followed by rapid hairpin refolding (Fig. 4a, light blue trace, *t* ≈ 10-12 s). As with unwinding, this pause occurs because of local duplex destabilization by the modification; the duplex below the modification cannot rezip until XPD has vacated enough nucleotides for an energetically favorable number of base pairs to reform (Fig. 4a, b top schematic, steps 2-4). However, for CPD, we also detected a pause at ∼37 bp (Fig. 4a, c, light arrow), preceding those at 35 bp. The pause at 37 bp suggests that damage-specific interactions must have occurred well past XPD’s front end. Its position is consistent with the internal pause detected during unwinding, ∼10-11 nt from the modification site, corresponding to the back end of XPD interacting with the modification. For fluorescein, the corresponding peak in residence times is shifted to ∼31 bp (Fig. 4c, light green arrow), again presumably due to linker stretching. Thus, the data suggest that many of the pauses observed with XPD unwinding are duplicated in this geometry, albeit in the reverse order due to the opposite XPD translocation direction (compare arrows in Fig. 3c and 4c).

We integrated the relative residence times over these regions to determine the excess traversal times (Fig. 4c, light shaded areas over 42-20 bp, Fig. 4d). XPD exhibited the greatest excess traversal time on the CPD site, τ*_exc_* = 2.9 ± 0.2 s, followed by the abasic site, fluorescein, and mismatch; τ*_exc_* = 0.7 ± 0.3 s, 0.5 ± 0.2 s, and 0.4 ± 0.2 s, respectively. With the exception of the internal pause for CPD (which we cannot measure during unwinding because a CPD rarely makes it into XPD‘s interior), the excess traversal times were consistently smaller than those exhibited during unwinding. These discrepancies may reflect differences in XPD interactions with the modifications due to the reversed fork-XPD geometry (see **Discussion**). Nevertheless, it is also possible that increased slips during rezipping on modified DNA could lead to underestimates of τ*_exc_*. **Tables 1, 2** provide a summary of all pause durations and positions.

We repeated this analysis on XPD rezipping substrates containing modifications on the 5’ strand, i.e. the displaced strand in this orientation. We found no appreciable effect of displaced strand modifications on XPD traversal probabilities nor on excess traversal times during rezipping (**Supplementary** Fig. 8).

## Discussion

XPD helicase is an essential component of the NER machinery and is considered to play a role in damage verification, though by mechanisms that remain unclear to-date. Our high precision, single-molecule measurements provide new insights into the detailed dynamics of damage sensing. We show that XPD is affected in distinct ways by different modifications, with CPDs—a natural substrate encountered in NER^4^—eliciting the strongest response. As XPD unwinds DNA, a single CPD site on the translocated strand blocks unwinding almost entirely (Fig. 2a). In comparison, an abasic site moderately decreases XPD’s likelihood of unwinding past the site, and a fluorescein modification has no impact on the level of subsequent unwinding. XPD’s ability to bypass a bulky adduct like fluorescein shows that damage sensing cannot occur purely by steric hindrance in or around XPD’s pore. This finding is consistent with recent structural^29^ and modeling studies^33,34^ that highlight the critical role of lesion-specific interactions.

XPD exhibits strong strand specificity, responding only to DNA modifications located on the translocated strand but never on the displaced strand, when either unwinding (Fig. 2a vs **Supplementary** Fig. 4c) or rezipping (**Supplementary** Fig. 7 vs **Supplementary** Fig. 8b). We note that this finding partially disagrees with previous biochemical studies on FacXPD^14^, which reported that helicase activity is inhibited by CPDs on either strand (albeit more strongly on the translocated strand), and with AFM imaging studies of taXPD^16^ and ctXPD^17^, which reported that different modifications are sensed on opposite strands. These discrepancies may result from differences between the XPD variants or the DNA substrate geometries used. As DNA is held under tension in our assay, it is possible that native contacts between the displaced strand and protein are disrupted. Furthermore, our assay geometry allows a DNA fork only at one end of XPD, which may preclude some native contacts present in other studies.

A previously unappreciated effect on the damage response is that of the XPD-fork geometry, highlighted in the differences in behavior observed during unwinding vs rezipping. Critically, while a CPD on the translocated strand blocks XPD during unwinding (Fig. 2a), it is bypassed efficiently during rezipping (**Supplementary** Fig. 7), during which the XPD-fork geometry is reversed from that of unwinding. One possibility to explain the difference in behavior is that the absence of a ss-dsDNA fork at the duplex separation point in XPD during rezipping alters the interactions made with the DNA lesion. Alternatively, it is possible that fork regression at the back end of XPD could provide a force to help push it through the damage site. Lending support to this model, we found that CPDs could be bypassed during unwinding under conditions in which multiple XPDs could bind and unwind DNA at the same time (**Supplementary** Fig. 9, see **Methods**). It was previously shown that multiple molecules of XPD could cooperate to increase unwinding processivity on undamaged DNA^41^.

Our measurements of XPD kinetics during encounters with DNA modifications provide additional details on damage sensing. Although each modification destabilizes the duplex, leading to rapid hairpin opening, the CPD, abasic, and fluorescein sites all slow down XPD unwinding, adding between 2 to 4 s to XPD’s mean transit time (Fig. 3d). Interestingly, fluorescein slows XPD’s progress the most during unwinding, which we suspect may be due to its bulk (see below). More importantly, we find no evidence that XPD forms a long-lived complex on a CPD, as has been proposed in multiple studies^14–17^. Instead, XPD most often backslides from the CPD site or dissociates completely in our measurements (Fig. 2b). Several seemingly disparate results may be reconciled given our observation that XPD can repeatedly shuttle between the CPD and hairpin base (Fig. 1d), which could be interpreted as a long stall at the lesion site at a lower spatial resolution. We also highlight that more direct single-molecule fluorescence measurements of FacXPD binding on CPD-containing DNA bubbles by Ghoneim et al.^18^ showed short XPD residence times of ∼7 s, more in line with our results.

The near base-pair resolution of our measurements of XPD dynamics on modified DNA allows us to infer that two regions of the helicase interact with the damage sites and to speculate on a structural basis for damage detection (Figure 5). For almost all modifications and XPD-fork orientations, we observe a primary ‘frontal’ pause when XPD’s front end presumably first encounters the DNA damage (shown schematically in Fig. 5c, d). The frontal pause locations during unwinding and rezipping (Fig. 3a, c and Fig. 4a, c, dark arrows) suggest that sensing occurs at or near the duplex separation point, located by the Arch, FeS and HD1 domains^29^ (Fig. 5a, right red circle). This picture is consistent with structures^29,31^ and computational models^33,34^ of XPD, which show a network of contacts made between DNA and these domains that may be involved in damage detection^33^, and with previous proposals of a lesion sensor pocket located near the entrance to the pore through which the translocated strand passes^15,27^. Figure 5b shows a model based on the human XPD structure (PDB 6ro4) highlighting residues in this sensing pocket (see **Supplementary Methods**; corresponding residues for XPD homologs shown in **Supplementary** Fig. 10). We speculate that the abasic site may similarly disrupt these contacts during unwinding, leading to a pause. Interestingly, we observe no pause for this modification in the rezipping configuration. Differences in damage sensitivity could reflect the different geometries of the DNA-XPD complex during unwinding vs rezipping. Again, we believe that the absence of a fork at the front end of XPD during rezipping could interfere with damage sensing interactions (Fig. 5d). For fluorescein, we attribute the positive shifts in pause locations for unwinding and negative shifts for rezipping to stretching of the linker on which the dye is attached. We thus propose that steric clashes between the bulky fluorescein and the small pore it must pass through may impede XPD’s progress. XPD’s remarkable ability to bypass fluorescein efficiently (Fig. 2a) suggests that the pore may need to open for translocation to resume. Honda et al.^43^ previously found that XPD can occasionally translocate over DNA-bound proteins without displacing them, illustrating an ability to negotiate even bulkier obstacles and lending support to the above mechanism.

**Figure 5.**
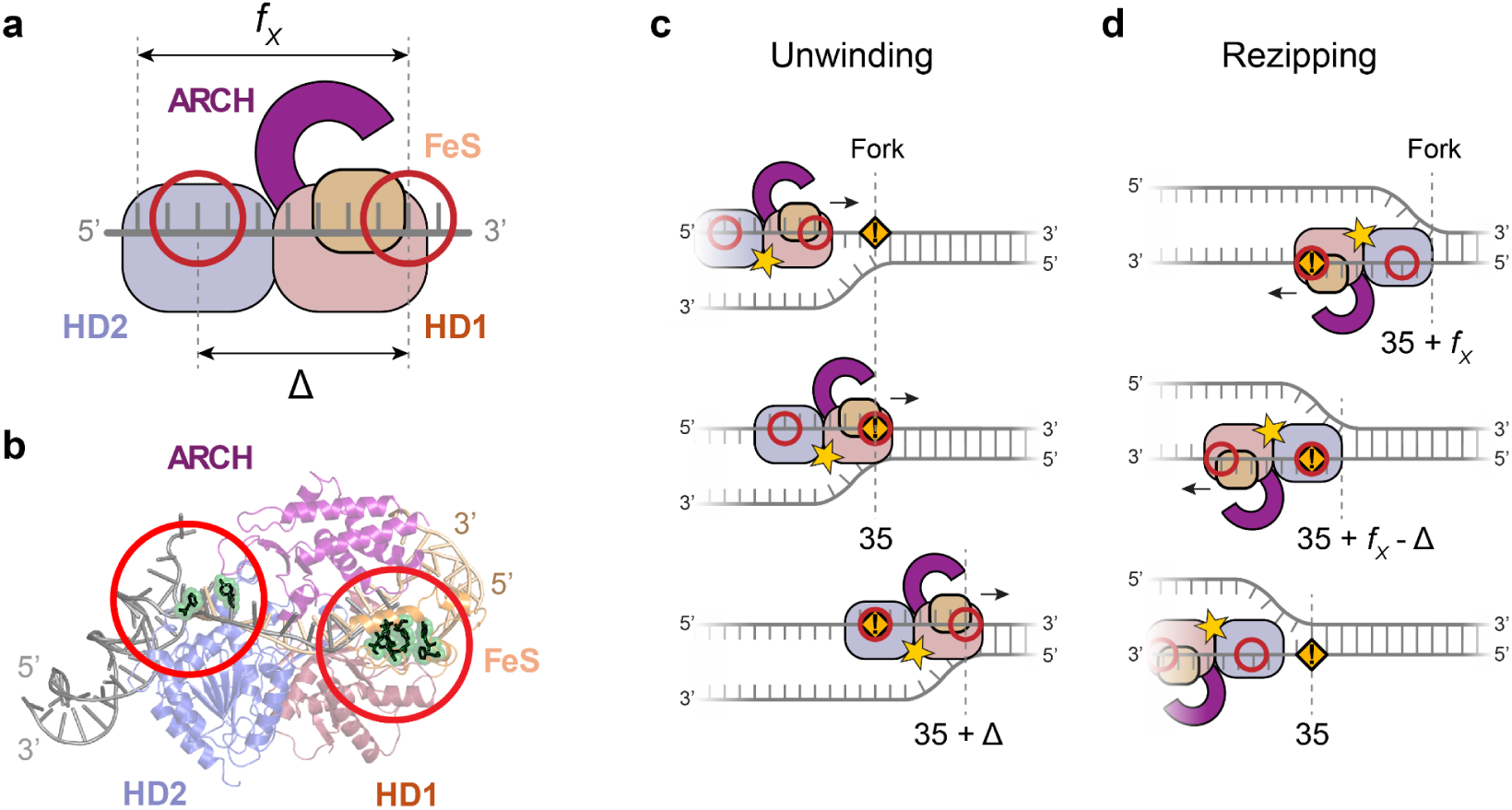
Model for XPD damage sensing. (**a**) Schematic of XPD and DNA with putative damage sensing sites (red circles). One damage sensing site is located near the ds-ssDNA fork separation point by the Arch, FeS, HD1 domains (right circle) and the other at a secondary interaction pocket on HD2 and by the ssDNA exit point (left circle). *f_X_*and Δ denote the footprint of XPD and distance between sensing sites, respectively. (**b**) Structural model of XPD (PDB 6ro4) with DNA. Two dsDNA-ssDNA fork structures are overlaid, one upstream of XPD (right, in tan color; PDB 8rev) at the duplex separation point during unwinding, and the other downstream of XPD (left, in gray color; PDB 6ro4), mimicking the proposed fork geometry during rezipping. The damage sensing sites corresponding to (a) are highlighted by the red circles, with residues proposed to be involved in damage detection displayed in green. (**c-d**) Schematics of XPD unwinding (c) and rezipping (d), and the DNA fork positions as the modification (at position 35) interacts with each damage sensing site (see also **Table 2**).

For most DNA modifications and geometries tested, we also see evidence for a second, ‘internal’ pause (Fig. 3a, c and Fig. 4a, c; light arrows), suggesting a secondary damage sensing site (see Fig. 5a, left red circle). Here, the pause positions collectively point to damage sensing ∼10-11 nt downstream of XPD’s front end (or ∼3 nt upstream from the exit point of the translocated strand), corresponding to the modification residing inside XPD’s interior on HD2 (see schematics in Fig. 5c, d). A number of structural studies have previously reported DNA contacts atypical for SF2B helicases in this region of XPD^27,30–32^ (**Supplementary Methods**). Interestingly, the location of this putative secondary damage sensing site is consistent with a region identified on HD2 of human XPD^31^, where ssDNA threads through a narrow passage lined with aromatic residues (highlighted in Fig. 5b, left red circle); this passage has been proposed to allow undamaged DNA to pass unimpeded but cause DNA lesions to pause^53^. Bulky adducts like fluorescein may also become blocked in this narrow channel, consistent with our observation of a secondary pause, shifted due to linker stretching (Fig. 3a**-c** and **4a-c**). The appearance of this internal pause for fluorescein modifications during both unwinding and rezipping suggests that these internal interactions are largely maintained in both XPD-fork geometries (Fig. 5c, d). CPD is an exception, although we suspect that this is simply because the internal pause is inaccessible during unwinding as a CPD rarely enters deeply into the XPD interior (Fig. 2a).

Figure 5 summarizes our model for both damage sensing sites. The schematics of XPD during unwinding (Fig. 5c) and rezipping (Fig. 5d) show the observed pause positions, marked by the DNA fork location when a modification interacts with frontal and internal damage sensing sites. **Table 2** shows that the measured pause positions for unwinding and rezipping compare well to predictions from the above model, given an XPD binding footprint *f_X_* = 14 nt, a distance between damage sensing sites Δ = 11 nt, and a fluorescein linker length *L_f_* = 7 nt. The structural model (Fig. 5b) overlays DNA forks at the front end (PDB 6ro4) and back end (PDB 8rev) of XPD, mimicking our proposed XPD-DNA configurations during unwinding (Fig. 5c) and rezipping (Fig. 5d), respectively.

Our findings shed light on the structural and kinetic mechanisms by which XPD may participate in DNA damage verification in the early stages of NER. Archaeal NER remains largely unclear, with the question of damage recognition still open, as there are no known homologs of the damage recognition protein XPC^54^. How DNA lesions are detected by XPD may thus provide important clues to this question. As XPD is part of a complex machinery in eukarya, an interesting question is how the above mechanisms could be impacted by XPD’s interactions with protein partners and vice versa. For example, we show that XPD’s response to DNA damage is more nuanced than simply stalling and locking onto the lesion. As others have noted^29,55^, the details of XPD’s interaction with damage may be important for positioning other components of the NER processing complex. XPG, responsible for the 3’ incision of the damaged fragment excised by NER, requires access to several nucleotides of ssDNA before the dsDNA junction to make an incision^56^. Thus, the prevalence of backsliding upon encounters with a CPD could be relevant in helping to clear the junction to grant access to downstream proteins. Studies of human NER factors show an interplay between different components in lesion detection^35,53,55^. Identifying how detailed interactions between XPD and other components of the NER machinery regulate damage sensing and excision is thus a rich subject ripe for future investigation.

## Methods

### Protein and DNA hairpin synthesis

*Fac*XPD was purified as described previously by Pugh et al.^39^. The hairpin constructs were synthesized using protocols adapted from Whitley et al.^45,46^. Briefly, the constructs consisted of two ∼1.5-kb dsDNA ‘handles’ (labeled ‘right’ and ‘left’) flanking an 86-bp ‘hairpin’ stem capped by a (dT)_4_ loop (Fig. 1b and **Supplementary** Fig. 1a**,c**). A 10-dT ssDNA binding site for loading a single molecule of XPD was positioned on the 5’ end of the hairpin stem. The ‘right handle’ (RH) and ‘left handle’ (LH) were synthesized by polymerase chain reaction (PCR) amplification of a section of lambda DNA (RH; New England Biolabs) and the pBR322 plasmid (LH; New England Biolabs), respectively. Primers functionalized with a single 5′ digoxigenin and biotin, respectively, were used during PCR for binding the final construct to anti-digoxigenin and streptavidin-coated beads. **Supplementary Table 1** provides the PCR primers used to synthesize RH and LH. The handles were digested with PspGI at 75 °C (LH) and TspRI at 65 °C (RH) for one hour. PspGI digestion generated a near-palindromic overhang. LH was dephosphorylated with antarctic phosphatase (37 °C for 30 min, 80 °C for 2 min) to minimize self-ligation. The hairpin, LH, and RH were then mixed in an equimolar ratio and ligated in a single step using T4 DNA ligase (22 °C for 1 h, inactivation at 65 °C for 15 min), and the final construct was gel-purified. DNA was purified with a QIAquick PCR purification kit (QIAGEN) after each reaction in the synthesis, except between LH digestion and dephosphorylation. A QIAquick Gel Extraction kit (QIAGEN) was used after gel purification. All enzymes were purchased from New England Biolabs.

We designed and synthesized two sets of hairpin stems for this study, with particular attention paid to sequence and stability. Since the unwinding dynamics of monomeric XPD is known to depend on the local base pairing stability of DNA^41^, a hairpin sequence with uniform stability throughout the stem presented a convenient platform for studying XPD’s response to DNA damage. We quantified the local stability of a hairpin sequence by calculating *P_open_*(*n*,*F*), the probability that DNA unwound to a given position, *n*, spontaneously opens further by one or more base pairs at a given tension *F. P_open_* is determined from the free energies of opening each base pair, which includes contributions from nearest neighbor interactions (including hydrogen bonding and base stacking)^47–49^ and from stretching the DNA to force *F*^41,49,57^. Two uniformly stable sequences were designed such that *P_open_* remained within a narrow range throughout the stem, up to base pair 75 (**Supplementary** Fig. 1b, d). From these two energetically uniform ‘core’ sequences, we constructed two sets of hairpin stems incorporating different DNA modifications: a 5’ set with the modification inserted on the side of the hairpin with the 5’ overhang (**Supplementary** Fig. 2a), and a 3’ set with the modification on the opposite, 3’ side (**Supplementary** Fig. 2d). Modifications were a single-base mismatch, a cyclobutane pyrimidine dimer (CPD), an abasic site, and a fluorescein conjugated to thymine via a 6-carbon linker, and were located at position 35, as measured from the stem base (or 34/35 for the 5’ CPD and 35/36 for the 3’ CPD). Two different mismatch substrates were synthesized for each set, with the mismatched base replacing either the native base at the modification position (labelled ‘on-strand’, **Supplementary** Fig. 2a, d), or the base on the opposite strand (’off-strand’, **Supplementary** Fig. 2a, d); these substrates were used to ensure that the translocated strand had the same sequence as that of the unmodified substrate during XPD unwinding and rezipping activity (i.e. the same 5’ strand for unwinding or 3’ strand for rezipping). **Supplementary Table 2** provides the full 200-nt core sequences of the hairpin inserts of the 5’ and 3’ set.

The hairpin stems with the core sequence were synthesized by self-annealing a single 200-nt oligonucleotide containing the sequence of interest and ligating it to the handles. This approach was not possible for hairpins containing the CPD modification, as oligonucleotides of that length with a CPD site were not commercially available. Instead, we synthesized hairpin stems incorporating modifications by annealing and ligating a set of ssDNA fragments. This piecewise design for the 5’ set comprised five fragments (**Supplementary** Fig. 1a): a ‘middle’ oligonucleotide on the 5’ strand in the center of the stem containing the modification (or native TT); the ‘middle complement’ oligonucleotide, providing a platform for the middle oligonucleotide to anneal and sticky ends to connect to the adjacent fragments; the LH and RH ‘base’ oligonucleotides, containing the base of the hairpin, the single-stranded protein binding site, and sticky ends for ligation to the handles and to the middle complement oligonucleotides; and the ‘loop’ oligonucleotide, forming a short hairpin with a dT-tetraloop cap and an overhang for annealing to the other side of the middle oligonucleotides. The 3’ sets of hairpins were constructed using a similar piecewise design (**Supplementary** Fig. 1c). To this end, we repurposed the modification-containing oligonucleotides from the 5’ set, placing them on the opposing strand of the stem, flipping much of the hairpin stem sequence. The other oligonucleotides were re-designed for the 3’ piecewise synthesis. In particular, since the LH base oligonucleotides became too short for stable hybridization, this base and the middle complement were combined into one ‘LH base + complement’ oligonucleotide. Additional details on the design and piecewise synthesis of the 5’ and 3’ sets of hairpins can be found in **Supplementary Methods**. Sequences of all oligonucleotides used for piecewise synthesis are given in **Supplementary Table 3**.

All oligonucleotides were purchased from Integrated DNA Technologies (Coralville, IA), except for the one containing a CPD, which was purchased from TriLink Biotechnologies (San Diego, CA). Where possible, all oligonucleotides, including primers, were ordered with PAGE purification.

### Optical trap

Previously described high-resolution dual-trap optical tweezers instruments^44–46^ were used to perform all the experiments. The traps were calibrated using standard procedures^44,58^. Data were acquired at rates of 100 Hz or 666 Hz (and downsampled to 95 Hz) using custom LabVIEW code.

XPD unwinding experiments were performed in buffer containing 100 mM Tris-HCl (pH ∼7.6), 20 mM NaCl, 3 mM MgCl_2_, 0.1 mg/mL bovine serum albumin (BSA; New England Biolabs) or recombinant albumin (rAlbumin, New England Biolabs). An oxygen scavenging system^59^ consisting of 2.9 U/mL pyranose oxidase (Sigma-Aldrich, St. Louis, MO), 0.15 mg/mL catalase (from *Aspergillus niger,* Sigma-Aldrich, St. Louis, MO), and 0.8% glucose was added to increase the lifetime of the DNA tethers^60^. To this buffer, XPD and ATP-γS or ATP were added, as described below.

### Sample flow chamber

Sample chambers consisted of a layer of parafilm (Nescofilm; Karlan, Phoenix, AZ) into which three channels were patterned, melted between two glass coverslips^46^. Samples and experimental buffers were flowed into the channels via syringe pumps (PHD 2000 Infusion or PHD Ultra; Harvard Apparatus, Holliston, MA; Nanojet, Chemyx, Stafford, TX). **Supplementary** Fig. 3 shows a schematic of the flow chamber. The upper channel (yellow) contained beads functionalized with anti-digoxigenin antibody (Adig beads); the lower channel (green) contained streptavidin-coated beads incubated with biotinylated DNA construct (SA-DNA beads). Beads were delivered through glass capillaries (OD = 100 μm, ID = 25 μm, Garner Glass Co., Claremont, CA) to the central channel (blue and red), where XPD unwinding activity was measured.

The central channel comprised two buffer streams merging under conditions of laminar flow^45,61^, which maintained a sharp interface between the bottom stream containing 500 µM ATP (red) and the top stream containing 20-22 nM XPD (with the exception of two traces at 5 nM) and 50 µM ATP-γS (blue). The spatial separation of XPD and ATP ensured that unwinding activity detected in the ATP stream arose from a single pre-loaded XPD molecule, as there were no other XPD molecules to bind to newly exposed ssDNA as the hairpin was unwound.

### Single-molecule unwinding assay

Prior to trapping, DNA was incubated for 1 h with streptavidin-coated beads (790 nm nominal diameter, Spherotech), then diluted with 100 mM Tris-HCl (pH ∼7.6), 20 mM NaCl, 3 mM MgCl_2_ for delivery to the sample chamber (**Supplementary** Fig. 3). Separately, an aliquot of protein G-coated beads (860 nm nominal diameter, Spherotech) functionalized with anti-digoxigenin antibody (Roche) was diluted for delivery to a different inlet of the sample chamber (**Supplementary** Fig. 3).

In a typical experiment, a bead of each type was trapped, and a single DNA tether was formed in the ATP stream. The quality of the hairpin was verified by mechanical unfolding and subsequent analysis of its force-extension curve (FEC). The DNA was then held at a constant tension of several pN, and the trapped, tethered beads were moved into the XPD stream. Once there, one molecule of XPD could bind to the 10-dT ssDNA loading site, but the absence of ATP in this stream precluded unwinding. After an incubation period of ∼1 min, the tension was raised to ∼12 pN, below the hairpin unfolding force, and the trapped, tethered beads were moved back into the ATP stream, where pre-loaded XPD could begin to unwind the hairpin. The absence of other molecules of XPD in the ATP stream ensured that any unwinding activity observed resulted from the single bound XPD molecule. Unwinding experiments described throughout the text were performed as described unless specified otherwise.

Some experiments (**Supplementary** Fig. 9) were designed to observe activity of multiple XPD molecules simultaneously. In these cases, the central channel consisted of the top stream containing both XPD and ATP and the bottom stream containing experimental buffer only. XPD binding to the ssDNA loading site could begin unwinding immediately due to the presence of ATP, and other XPD molecules in solution could also bind onto the released ssDNA and potentially interact with XPD:DNA complexes. In these experiments, multiple turnovers of XPD activity were expected since both XPD and ATP were present and constantly replenished by flow through the chamber.

## Data analysis

### Analysis of force-extension curves

Prior to each measurement of XPD unwinding, a FEC of the DNA molecule tethered between the two trapped beads was collected by changing the trap separation at a constant rate of 100 nm/s. The FECs displayed a characteristic transition at ∼15 pN when the hairpin stem mechanically unfolded (**Supplementary** Fig. 2b, e). Sections of the FECs above and below the transition force were fit to models of the DNA handles and of the handles plus unfolded hairpin, respectively (dashed lines), based on the extensible worm-like chain (XWLC)^62^. Parameters for ssDNA and dsDNA elasticity were obtained to provide the best fits to the FECs and were consistent with previous studies^63–65:^ persistence length, *P_ds_* = 50 nm and *P_ss_* = 1.07 nm, inter-phosphate distance, *h_ds_* = 0.34 nm/bp and *h_ss_*= 0.6 nm/nt, and stretch modulus *S_ds_* = 1000 pN and *S_ss_*= 1000 pN. Agreement with the models and visual inspection of the unfolding transitions confirmed that the hairpin was properly synthesized.

To compare the destabilizing effect of each modification, we determined the mean and standard deviation of selected FECs for each construct over 3-nm extension bins (**Supplementary** Fig. 2b, e, solid lines and shaded areas, respectively). We then aligned the unfolding transitions of the average FECs for the 5’ and 3’ set separately and integrated the areas under the transitions (limits of integration indicated by the dashed blue lines). Finally, we calculated the differences between the area under the transition for each construct containing a modification and that for unmodified DNA (**Supplementary** Fig. 2b, e, insets). These areas represent the mechanical work to unfold modified DNA relative to that to unfold unmodified DNA, and provide a measure of the destabilizing effect of each modification. Error bars denote the standard deviation and were determined by summing in quadrature the errors of the mean FECs at individual extensions.

### Selection and analysis of hairpin unwinding and rezipping trajectories

XPD unwinding traces were collected at a constant tension (of mean ∼12 pN, below the hairpin unfolding transition) using active force feedback. To calculate the number of base pairs unwound over time during XPD activity, the change in extension (in nm) of the DNA construct was divided by the extension of the two nucleotides of ssDNA released at the measured force, using the XWLC parameters for ssDNA referenced above.

A selection of the single-molecule trajectories collected were analyzed based on low noise characteristics and stability in baseline, determined from shifts in the extension of the DNA tether prior to unwinding and after XPD dissociation. Within each trajectory, we analyzed only those bursts for which XPD reached or exceeded 25 bp. Restricting the analysis to high-processivity bursts enabled a more consistent comparison of XPD activity. Bursts were further split into segments before and after the burst peak, corresponding to unwinding and rezipping, respectively. In cases where the peak contained a plateau or some minor rezipping (≲6-8 bp) followed by unwinding to within 3-4 bp of the burst maximum, these segments were included in the unwinding portion, as we considered XPD less likely to be on the 3’ strand than the 5’ strand in these instances. **Supplementary Table 4** lists the numbers of bursts analyzed under each set of conditions.

### Calculation of traversal probability and errors

To determine the probability that XPD could bypass a modification during unwinding, we first determined the maximum position reached by each burst and then counted the total number of bursts, *n*, that remained at each base pair position on the hairpin for a particular DNA substrate. The traversal probabilities were then calculated using the Laplace estimator *p* = (*n* + 1)/(*N* + 2), where *N* was the total number of bursts that survived to 35 bp on that substrate. (The Laplace estimator provides a less biased estimate of the success probability than *n*/*N*.) The errors were determined from the expression for the variance of the Laplace estimator σ^2^ = (*n* + 1)(*N* - *n* + 1)/(*N* + 2)^2^(*N* + 3).

To determine traversal probability during rezipping, we restricted our analysis to XPD translocating down the 3’ strand specifically. We analyzed only bursts displaying gradual (*bona fide*) rezipping at 50 bp, when XPD would first come into contact with the modification while translocating down the 3’ strand. Bursts with a backslide through 50 bp were excluded from the analysis as XPD could conceivably be sliding down either strand. Successful traversal was defined as continued gradual rezipping through the region around the modification, without dissociation or backslide (identified from ≥10 bp drops). The traversal probabilities and errors were then calculated using the formulas above.

### Residence time analysis

For each unwinding or rezipping segment of a burst, we defined the residence time at a given position as the total amount of time that the DNA fork resided at that position. We note that the residence time as defined here could arise from multiple excursions to a particular position from surrounding positions during a single burst, and is distinct from the dwells during XPD stepping as described in ref. ^41^. For each individual unwinding or rezipping segment, the residence time at each position on the hairpin was determined by kernel density estimation (KDE) using a Gaussian kernel with a width σσ = 1 bp. Bursts did not contribute residence times at positions they did not reach. The mean and s.e.m. in residence times were then determined from the set of unwinding or rezipping segments analyzed. Residence times at each position for the different hairpin constructs with 5’ modifications are shown in **Supplementary** Fig. 5. To compare XPD dynamics on modified DNA to those on the reference, core construct, we calculated the difference between the mean respective residence times—or the relative residence time—at each position (Fig. 3c and 4c). Errors were calculated by adding the errors of the mean residence times on the modified and unmodified DNA in quadrature.

To determine how long XPD took to traverse the damage-containing DNA—or the excess traversal time—we integrated the relative residence times over a range of positions, as indicated by the shaded areas in Fig. 3c and 4c. Fig. 3a and 4a show that the modifications destabilized the duplex, leading to spontaneous fork opening and a pause at the modification site. To account for this effect in determining the excess traversal times, we integrated over the destabilized region through the modification site (as indicated in each figure). Errors for the excess traversal time were determined by summing in quadrature the errors of the mean relative residence times at individual positions.

For rezipping, residence time analysis was restricted to segments demonstrating gradual (*bona fide*) rezipping, indicating that XPD was translocating down the 3’ strand. As backslides could presumably occur from either strand, they were only included if preceded and followed by clear *bona fide* rezipping, indicating that XPD remained on the 3’ strand throughout.

Bootstrapping analysis was used to identify outliers in residence time distributions, for unwinding and rezipping activity on each modification. Multimodality in the bootstrapped distributions at a given fork position indicated that a burst contributing an unusually long residence time at that position was present in the sample. Considering residence times around the modification only (20-50 bp), we identified and removed outlier bursts in their entirety from the residence time analysis to produce more consistent mean dwell times. In total, this procedure resulted in the removal of two mismatch bursts for 5’ rezipping; two fluorescein bursts for 3’ set unwinding; and one burst each of unmodified, abasic, fluorescein, and mismatch for 3’ set rezipping.

## Data availability

The data that support the findings of this study are available from the corresponding author upon reasonable request.

## Code availability

All experimental data were collected from custom-built optical tweezers instruments operated with custom LABVIEW code. Analysis of experimental data was carried out using custom MATLAB code. Code is available from the corresponding author upon reasonable request.

## Acknowledgements

We thank members of the Chemla and Spies labs for helpful discussions. This work was supported by National Institutes of Health grants R35 GM144125 (to Y.R.C.) and R01 CA232425 (to M.S.).

