## Supplementary Information for "Base-pair scale dynamics of a repair helicase on DNA lesions reveal varied damage-sensing mechanisms"

Table of contents:

Supplementary Methods: p. 1-5

Supplementary Figures 1-10: p. 6-15

Supplementary Tables 1-4: p. 16-18

Supplementary References: p. 19-20

### Supplementary Methods

#### Construct design for hairpins containing modifications

As outlined in **Methods**, constructs with the core (unmodified) hairpin sequence were made by ligating the left and right dsDNA handles to a single 200-nt oligonucleotide. However, CPD-containing oligonucleotides of 200-nt length were not commercially available. We therefore synthesized hairpin stems by ligating together multiple shorter fragments: an oligonucleotide containing a CPD (or other modification), and a set of unmodified oligonucleotides (**Supplementary Fig. 1a, c**, **Supplementary Table 3**). This piecewise design gave us the flexibility to make a range of hairpin constructs with the same core sequence but different modifications. Our approach was similar to a scarless Golden Gate assembly[1], without cleavage by restriction enzymes: each hairpin stem consisted of annealed pairs of complementary ssDNA fragments with overhangs, attached together in a unique order through single-step ligation. In the end, the 5' set and 3' set of hairpins were composed of 5 and 4 oligonucleotides respectively, between 22 and 87 nt long.

The choice of ssDNA fragments for piecewise assembly involved multiple considerations. First, fragments needed to hybridize efficiently and stably to their complement, which depended on their lengths and sequences. As our energetically uniform core sequence is repetitive, our design had to account for alternate secondary structures which could interfere with assembly. Also, since the modifications assayed destabilize the surrounding duplex[2,3], our design had to be robust to lower stability in these regions. Second, the overhangs (or sticky ends) needed to be chosen for each fragment to ligate rapidly and with high fidelity to its correct neighbor. The efficiency and fidelity of ligation by T4 ligase can vary drastically depending on the overhang sequence[4,5]. We relied heavily on UNAFold[6–8] and DINAMelt[9] to assess hybridization and secondary structure of prospective fragments and on the comprehensive studies of sequence-dependent ligation efficiency by Potapov et al.[4,5] to design breakpoints in the core sequence yielding overhangs that would ligate well. The high cost of CPD-containing oligonucleotides placed an additional constraint on the fragment and breakpoint designs. The CPD-containing ssDNA fragment ideally needed to be as short as possible, without compromising hybridization to its complement, for maximum reusability in different pre-existing hairpin sequences. Consequently, we selected a 27-nt oligonucleotide with a CPD located 4 nt from the 5' end, as explained below.

The piecewise design for the 5′ set of constructs is shown in **Supplementary Fig. 1a** and the oligonucleotides used are listed in **Supplementary Table 3**. To determine the length of the 'middle' fragment that contained the DNA modification, we used DINAMelt to estimate the hybridization efficiencies of different potential lengths of 'middle':'middle complement' oligonucleotides. Since

DINAMelt could not model a CPD, abasic, or fluorescein site explicitly, we mimicked the duplex destabilizing effect of a modification by introducing a single-base mismatch in the modeled 'middle' oligonucleotide. The junction between the 'loop' and 'middle' oligonucleotides consisted of a 3-nt, 5' sticky end, selected by applying the comprehensive ligation efficiency data of Potapov et al.[4,5]. Using these tools, we compared the number of total ligations, fractions of correct ligations, and the fragments that mis-ligated for different possible lengths of 'middle' oligonucleotide with 3-nt overhangs. Of the possible choices, a 27-mer appeared optimal, giving exceptionally high fidelity and relatively high total ligation. For the junction between the 'base' oligonucleotides and the 'middle':'middle complement' duplex, we opted for a 3' overhang, 9 nt in length. Since the modification was only 4 nt from the edge of its fragment due to reusability constraints, we reasoned that a 5' sticky end would leave too short a hybridization region and that the destabilizing effect of the modification would interfere with ligation. Instead, a 3' 9-nt overhang allowed us to move the junction away from the modification and provide a sufficiently long hybridization region for efficient ligation to the 'base' oligonucleotides. Finally, the choice of breakpoints at the ends of the 'middle':'middle complement' oligonucleotides determined the 'base' and 'loop' fragments. Overall, DINAMelt predicted that designed fragment pairs would hybridize efficiently at room temperature. However, it also predicted the presence of significant secondary structures in some fragments. During synthesis, we tempered the effect of alternate structures by slowly annealing some oligonucleotides, allowing them to form the most stable fully hybridized structure.

We used a similar approach to design the 3' set of constructs (**Supplementary Fig. 1c** and **Supplementary Table 3**). Here, the 'middle' oligonucleotide containing the modification was re-used, but now flipped to the opposite strand of the hairpin stem, with the modification at 35 bp (35/36 bp for the CPD) from the hairpin base. To retain maximum sequence context, we also flipped as much of the hairpin sequence surrounding the 'middle' oligonucleotide as possible. Consequently, the 5' → 3' core sequence over positions 1-69 on the 5' strand of the 5' set of hairpins is identical to that on the 3' strand of the 3' set (compare black arrows in **Supplementary Fig. 1a** and **c**). As a consequence, XPD encounters the same sequence on the translocating strand while rezipping on the 3' set as it does unwinding on the 5' set over this range. We note that $P_{open}$ for the sequence of this 3' set fluctuated beyond the range of the 5' set (compare **Supplementary Fig. 1b** and **d**), but this excursion took place in the 'middle' oligonucleotide region, which could not be modified. One difference between the 3' and 5' sets involved the 'middle complement' fragment design. Since the position of the 'middle' oligonucleotide in the 3' set of constructs provided a short hybridization region with the 'base' oligonucleotides, we opted to combine the 'middle complement' fragment with the adjacent 'base' adapter to form an extended complement ('base + middle complement') that would be more stably hybridized.

### Piecewise synthesis for hairpins containing modifications

For the synthesis of hairpins in the 5' set, the 'middle' and 'middle complement' oligonucleotides were each diluted to 10 µM with nuclease-free duplex buffer from Integrated DNA Technologies (30 mM HEPES, pH 7.5; 100 mM potassium acetate). Other oligonucleotides were each diluted to 10 µM with IDTE (10 mM Tris, 0.1 mM EDTA, pH 8; Integrated DNA Technologies). Equimolar amounts of the 'middle' and 'middle complement' oligonucleotides were combined and annealed slowly in a thermal cycler as follows: at 95 °C for 5 minutes, then decremented by 0.5 °C/min, using a 0.1 °C/s ramping rate, to 10 °C. The resulting duplex and the 'base LH', 'base RH', and 'loop' oligonucleotides were mixed in equimolar amounts and ligated with T4 DNA ligase (New England Biolabs) at room temperature for 3-5 hours. Purifying fully-ligated hairpin from unligated fragments was not practical due to the short total length. Instead, based on previous characterization, we considered that the reaction had gone to completion to calculate molar amounts for the final full-construct ligation with handles. While some unligated fragments remained in the hairpin ligation mixture, we obtained yields of the final constructs between 20 and 70 nM, sufficient for our single-molecule studies.

For the 3' set, oligonucleotides were separately diluted to 10 µM with nuclease-free duplex buffer. The full set of oligonucleotides for synthesis were combined in equimolar amounts, then annealed slowly and ligated with T4 ligase as described above. Yields of the final constructs were between 15 and 75 nM.

### Structural analysis of XPD and damage sensing sites

To interpret the dynamics observed in our single-molecule measurements, we analyzed XPD structures for insights on XPD's interactions with DNA and potential damage sensing sites. Using PyMOL, we compared three structures of XPD homologs that feature interactions with DNA: (i) *Chaetomium thermophilum* (ct)XPD with its protein partners (PDB entry 8rev)[10] showing the helicase on a cross-linked ds-ssDNA fork (**Supplementary Fig. 10a**); (ii) the human TFIIH core with XPA (PDB entry 6ro4)[11] showing human (h)XPD on a ds-ssDNA fork located at the tail of the helicase by HD2 (**Supplementary Fig. 10b**); and (iii) the archaeal *Thermoplasma acidophilum* (ta)XPD with a small fragment of ssDNA bound to HD2 (PDB entry 4a15; **Supplementary Fig. 10c**)[12]. Since no FacXPD structures have been reported to-date, we used multiple sequence alignment (analyzed using Clustal 2.1[13]; data not shown) to speculate on DNA interactions in *F. acidarmanus*. **Figure 5b** displays the hXPD structure with its bound DNA (PDB 6ro4) and the DNA fork from the ctXPD structure (PDB 8rev) overlaid, and provides a model for the fork geometries during both unwinding and rezipping activities. We used this combined structure to enumerate the nucleotides between forks and estimate XPD's binding footprint on ssDNA, $f_X \approx 14$ nt (see **Table 2**).

We also used these structures to locate potential damage sensing sites. Multiple residues on HD1/FeS near the entry pore for ssDNA are implicated in damage sensitivity (highlighted in green on HD1/FeS in **Supplementary Fig. 10a-c**). In hXPD, the network of Y158, F161, and F193 on FeS have been proposed to proofread DNA bases (**Supplementary Fig. 10b**)[11]. In addition, R112 and C134 bridge DNA to the FeS cluster[11], which could be involved in DNA damage detection through charge transfer[14,15], and Y192 and R196 bind to the DNA phosphate backbone near the pore in the structure[11]. The latter two residues are highly conserved across XPD homologs and have been shown to impair DNA damage sensing when mutated[16]. In ctXPD, the homologous Y191 and R195 similarly interact with the DNA phosphate backbone (**Supplementary Fig. 10a**). The corresponding residues in taXPD are Y166 and K170 (**Supplementary Fig. 10c**) and in FacXPD Y171 and R175[16], respectively. K170 mutants are known to lose sensitivity to DNA lesions[17]. Interestingly, K170 is not conserved in *Sulfolobus acidocaldarius* (Sac)XPD, which may explain why SacXPD shows little sensitivity to damage[18,19].

On HD2 of human XPD, F508 and Y627 (green residues on HD2, **Supplementary Fig. 10b**) form base-stacking interactions with DNA near the ds-ssDNA junction in the structure[11], unusual as SF2 helicases canonically interact with the DNA backbone. These residues may be good candidates for sensing damaged bases. Indeed, Kim et al.[20] proposed that aromatic residues (including F508 and Y627) lining the narrow passage through which ssDNA is threaded may allow normal ssDNA to pass but block DNA lesions, potentially forming a site for damage sensing on HD2 a distance $\Delta \approx 11$ nt from the entry pore (see **Table 2**). Since the overall orientation and DNA interactions are highly comparable in the ctXPD structure, the corresponding HD2 residues, F507 and Y626, can be inferred to interact similarly with ssDNA (**Supplementary Fig. 10a**). In taXPD, Y425 and R529 are the corresponding residues in HD2 identified by our sequence alignment (**Supplementary Fig. 10c**). A structure of taXPD[21] (PDB 5H8W; not shown in **Supplementary Fig. 10**) resolves 4 nt of ssDNA bound to HD2 and shows both of these residues stacking with DNA bases, suggesting similar ssDNA interactions. Additional base-stacking interactions are also seen in HD2 of taXPD with F538 and P530 in the same structure (PDB 5H8W) and with W549 in a separate structure (PDB 4a15)[12]. The prevalence of stacking interactions in the same region of XPD may indicate significant functional relevance, and the direct interactions with DNA bases suggest a possible role in damage sensing.

### Supplementary Figures

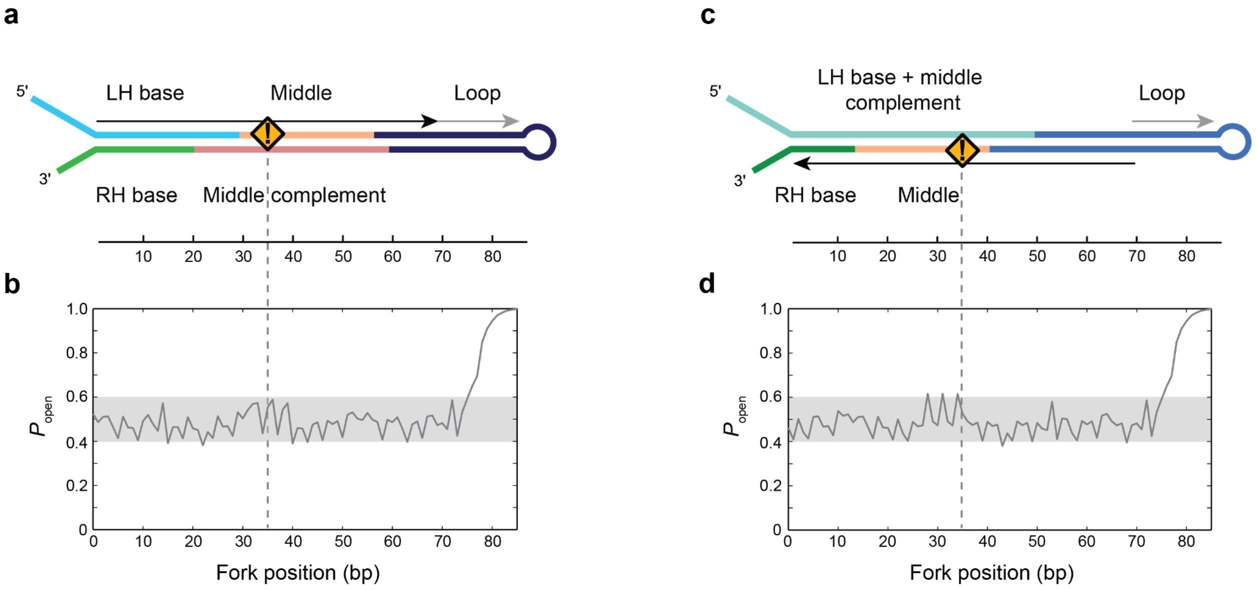


**Supplementary Figure 1 | Hairpin constructs design.** (**a**) Schematic for the piecewise design of the hairpin constructs containing a 5’ strand modification at position 35 bp (orange warning sign). Oligonucleotides are annealed and ligated to form the 86-bp hairpin stem, which is then ligated to ~1.5-kb DNA handles (see **Methods**). (**b**) $P_{open}$ vs hairpin position for the core sequence into which all 5’ modifications are inserted. The core sequence is designed such that $P_{open}$ is uniform within a narrow range (shaded area). (**c-d**) Same as a-b, for constructs containing a 3’ strand modification. Black and gray arrows in a and c indicate sequences that are identical in the two constructs.

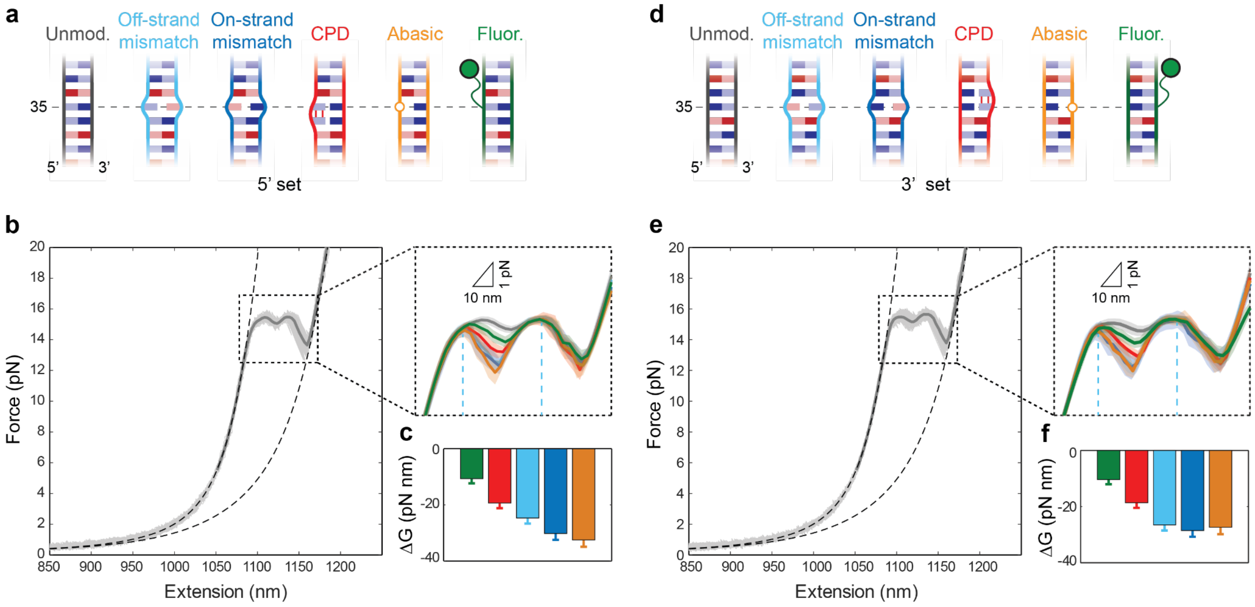


**Supplementary Figure 2 | Hairpin force-extension behavior.** (**a**) Schematics of the constructs containing a 5' strand modification, showing the unmodified core sequence (gray), off-strand single-base mismatch (light blue), on-strand mismatch (blue), cyclobutane pyrimidine dimer (CPD; red), abasic site (orange), and fluorescein (green). (**b**) Representative force-extension curves (FEC) for the unmodified core sequence (gray) for the 5' set. Dashed lines indicate theoretical models of the folded and unfolded construct FECs. Inset shows hairpin unfolding transitions for the 5' set of constructs, manually aligned to overlap (same colors as a; dark lines denote the average FECs and the shaded areas the standard deviation). (**c**) Integrated area under the unfolding transition for each construct (same colors as a) relative to that of the uniform core sequence, representing the destabilization of the duplex by each modification. Blue dashed lines indicate the integration range. (**d-f**) Same as a-c, for constructs containing a 3' strand modification.

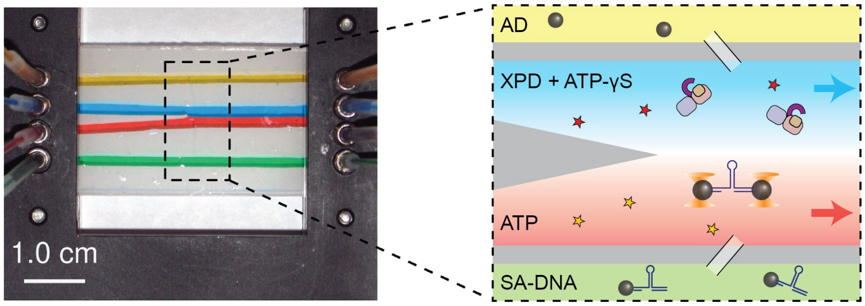


**Supplementary Figure 3 | Laminar flow cell.** Sample chamber layout containing 4 inlet and outlet streams (left, photograph; right, schematic, not drawn to scale). Two adjacent streams containing XPD + γS-ATP (blue channel) and ATP only (red), respectively, merge into a central channel but do not mix appreciably due to the laminar flow. During an experiment, a tether is incubated in the protein stream, and then moved across the stream interface and into the ATP-containing stream to initiate unwinding. The top/bottom channels (yellow/green) are loaded with anti-digoxigenin (AD) beads and streptavidin beads with attached DNA (SA-DNA), respectively. The two types of beads diffuse into the central channel via thin glass capillaries. Photograph copyright Qi et al.[22], distributed under the terms of the Creative Commons Attribution License (https://creativecommons.org/licenses/by/3.0/).

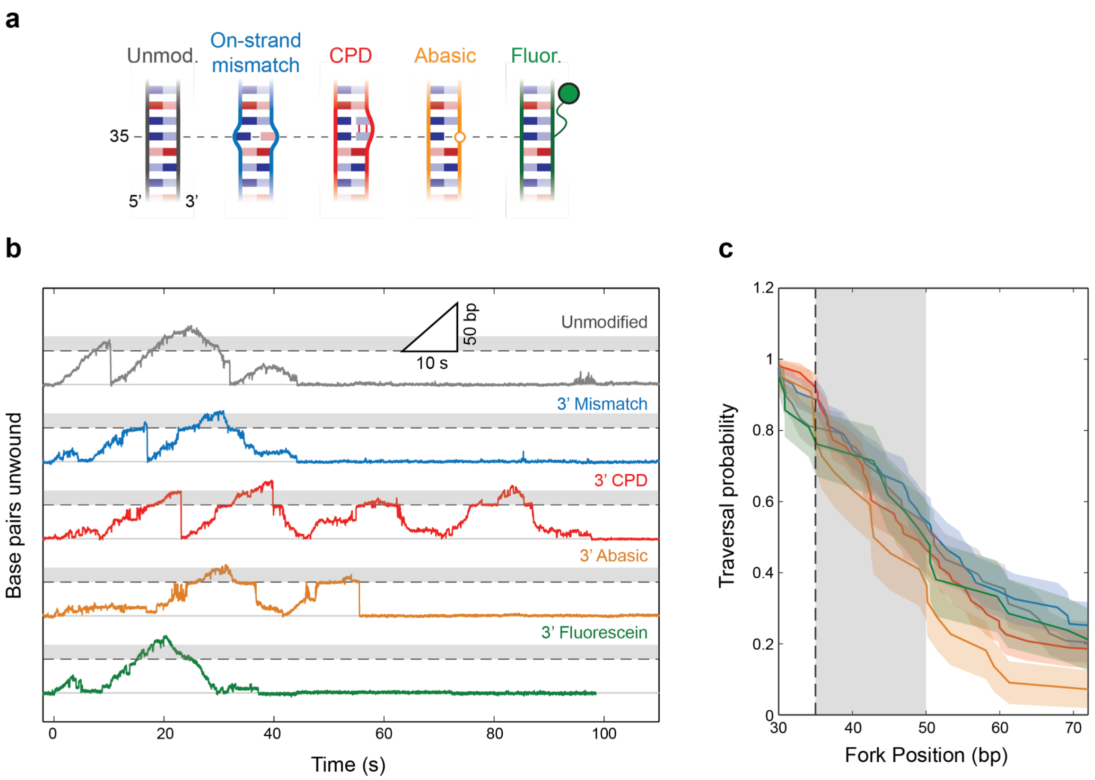


**Supplementary Figure 4 | XPD unwinding on 3' displaced strand modifications.** (**a**) Schematics of the DNA substrates with 3' strand modifications: unmodified 'core' sequence (gray), single-base mismatch (blue). CPD (red), abasic site (orange), and fluorescein (green). (**b**) Representative unwinding traces on each DNA construct (same colors as a). Traces are offset for clarity; the dashed lines indicate the positions of the base of the hairpin and the modification. The shaded region between 35 and 50 bp indicates the region where the modification interacts with XPD. (**c**) Traversal probabilities past the modification for each DNA construct (same colors as a). Shaded regions denote standard error of the mean (s.e.m.).

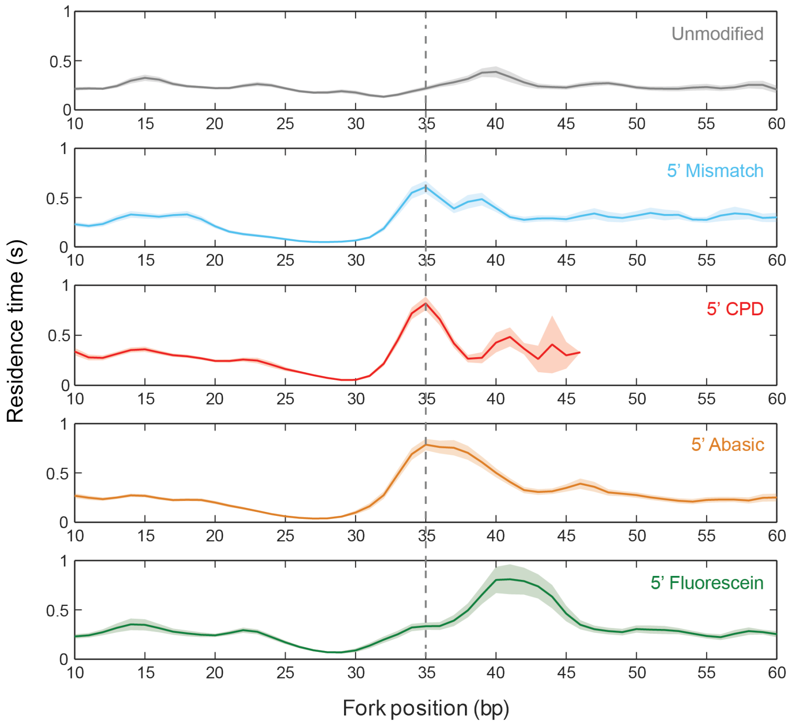


**Supplementary Figure 5 | XPD residence time on 5'-strand modifications.** Average residence time vs DNA fork position on each substrate: unmodified core sequence (gray), single-base mismatch (light blue), CPD (red), abasic site (orange), and fluorescein (green). Shaded areas denote s.e.m..

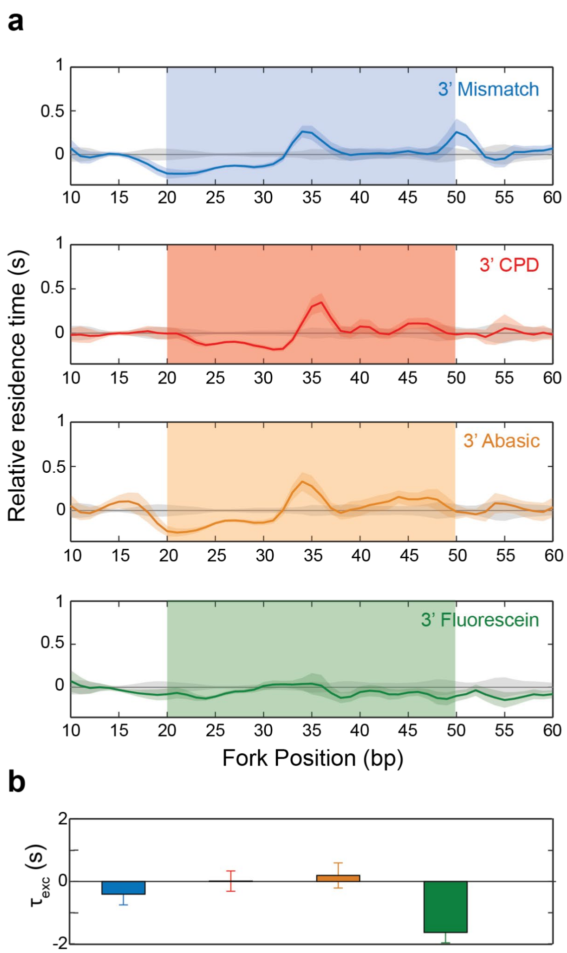


**Supplementary Figure 6 | XPD unwinding residence times on 3' displaced strand modifications.** (**a**) Average residence time vs DNA fork position on each modified substrate (same colors as a), measured relative to that on unmodified DNA (gray). Shaded areas denote s.e.m. (**e**) Excess XPD traversal times, $\tau_{exc}$, for different substrates (same colors as a). (Note: negative values of $\tau_{exc}$ may arise from the difference between XPD's unwinding speed and its translocation rate, which would cause it to catch up to the fork ahead faster than it would unwind through the same length of unmodified dsDNA.) Error bars denote s.e.m..

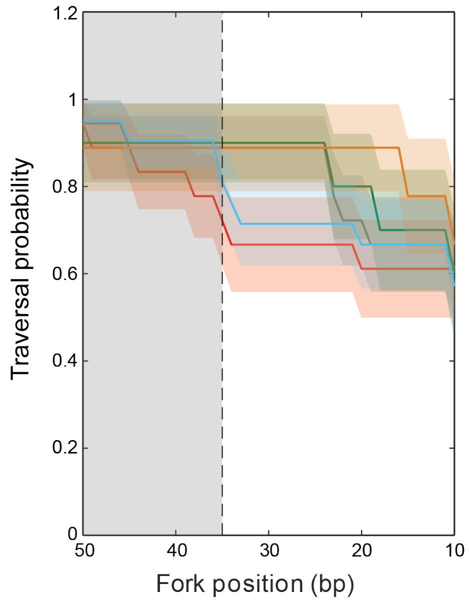


**Supplementary Figure 7 | Traversal probabilities for rezipping on 3'-strand modifications.** Traversal probabilities for XPD rezipping past the modification for each DNA construct (same colors as Figure 1). The modification site and interaction region are denoted by the dashed line and gray shaded area.

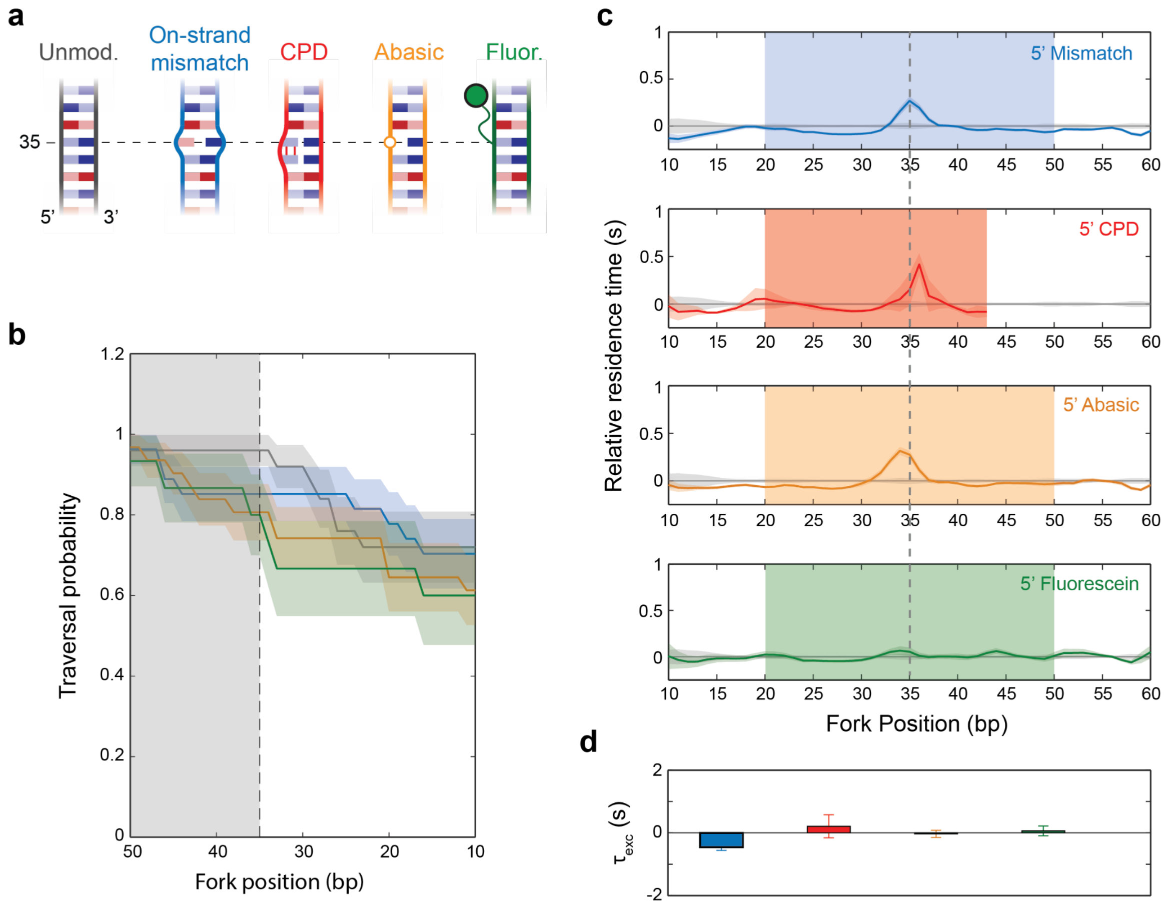


**Supplementary Figure 8 | XPD rezipping on 5'-strand modifications.** (a) Schematics of the DNA substrates with 5' strand modifications: unmodified 'core' sequence (gray), single-base mismatch on the translocated strand (blue). CPD (red), abasic site (orange), and fluorescein (green). (b) Traversal probabilities past the modification for each DNA construct (same colors as a). Shaded regions denote s.e.m. (c) Average residence time vs DNA fork position on each modified substrate (same colors as a), measured relative to that on unmodified DNA (gray). Shaded areas denote standard deviation. (d) Excess XPD traversal times, τexc, for different substrates (same colors as a). Error bars denote s.e.m..

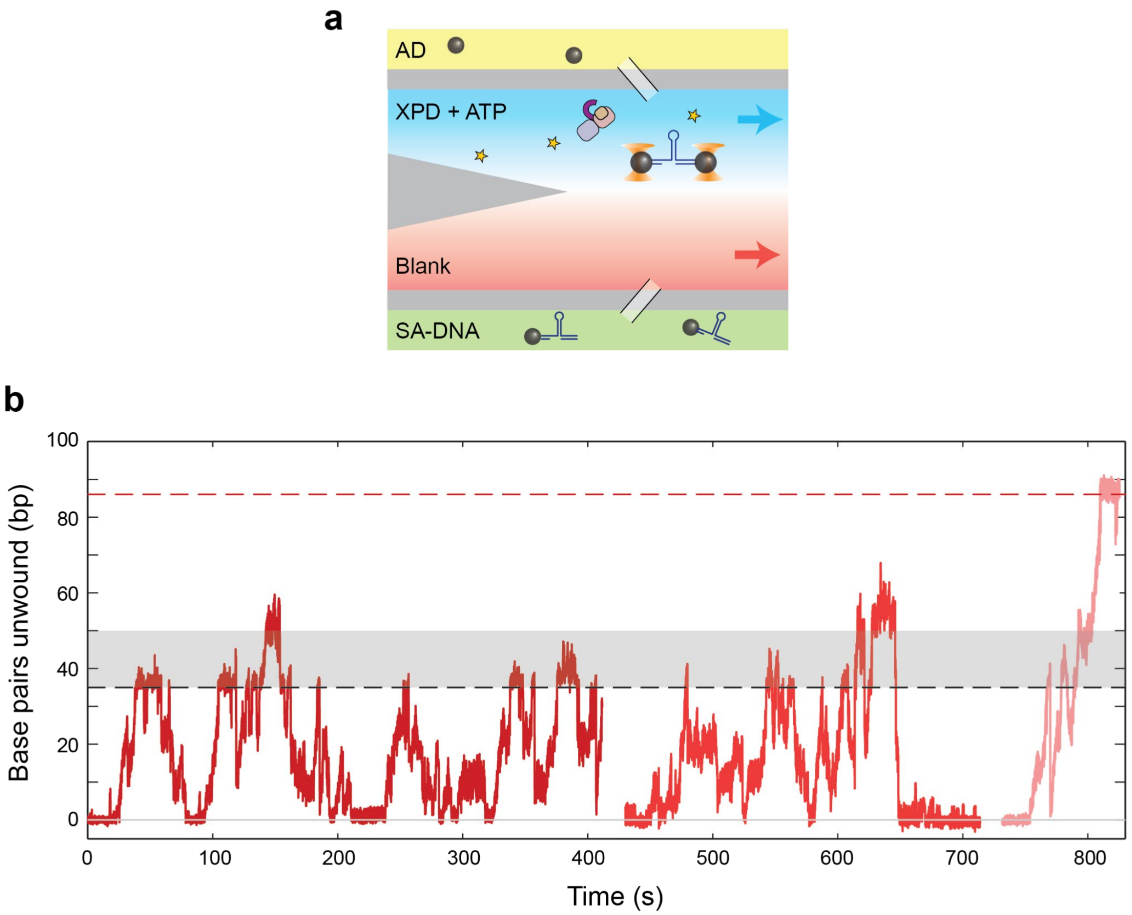


**Supplementary Figure 9 | Multiple XPD can collectively bypass a 5'-strand CPD.** (**a**) Laminar flow cell for monitoring the activity of multiple XPD molecules simultaneously (see SI Fig. 1 for comparison). Here, the two inner streams contain XPD + ATP (blue channel) and buffer only (red). During an experiment, a tether is incubated in the blank stream, then moved into the ATP-containing stream to initiate unwinding. Other XPD molecules in solution can bind onto the released ssDNA. (**b**) Example traces of multiple XPD's unwinding a hairpin containing a 5'-strand CPD, showing several instances where the CPD site (position 35 bp; black dashed line) is bypassed.

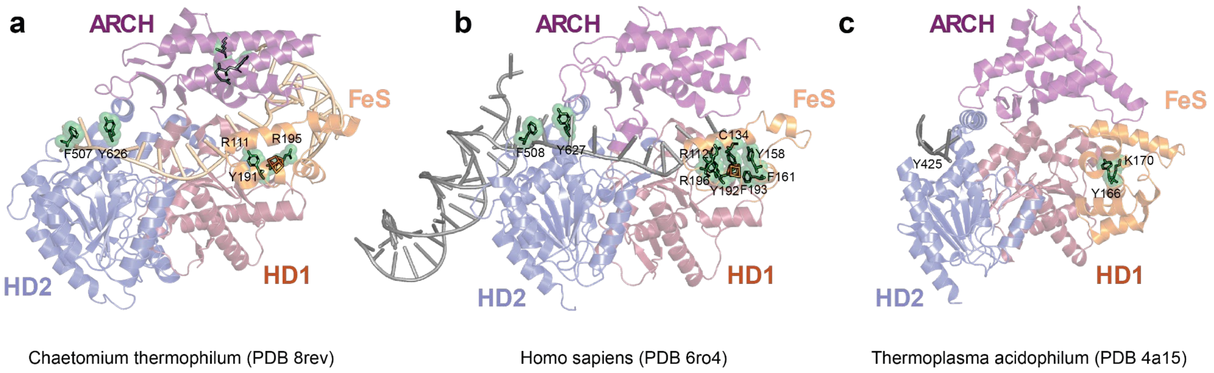


**Supplementary Figure 10 | Comparison of XPD variants and putative sensing sites.** (**a-c**) Structures of three XPD variants in complex with DNA and with residues proposed in damage detection highlighted (green): (a) *Chaetomium thermophilum* (ct)XPD structure with ds-ssDNA fork (PDB: 8rev), (b) human XPD structure with ds-ssDNA fork (PDB: 6ro4), and (c) *Thermoplasma acidophilum* (ta)XPD structure (PDB: 4a15) with bound ssDNA fragment.

### Supplementary Tables

| Primer name | Sequence (5'-3') |
|---|---|
| RH forward primer | /5DigN/ GGG CAA ACC AAG ACA GCT AA |
| RH reverse primer | CGT TTT CCC GAA AAG CCA GAA |
| LH reverse primer | CAA GCC TAT GCC TAC AGC AT |
| LH forward primer | /5Biosg/ TGA AGT GGT GGC CTA ACT ACG |

**Supplementary Table 1 | Primers used for synthesis of dsDNA right handles (RH) and left handles (LH).** Sequences are shown in nomenclature of Integrated DNA Technologies (IDT). Abbreviations correspond to the following modifications: DigN = digoxigenin and Biosg = biotin.

| Oligonucleotide name | Sequence (5'-3') |
|---|---|
| 5' set core sequence | /5Phos/CC TGG TTT TTT TTT T AG TCT CAG TCA CTC ATG TCA GTC ACA GTC AGT CTT GAT GAT GTC ACT GAC TGA GAC TCT GAC TCA CTG AGT CAT GTC TGA GAG TCG TTT TCG ACT CTC AGA CAT GAC TCA GTG AGT CAG AGT CTC AGT CAG TGA CAT CAT CAA GAC TGA CTG TGA CTG ACA TGA GTG ACT GAG ACT CCC ACT GGC |
| 3' set core sequence | /5Phos/CC TGG TTT TTT TTT TTC AGT GAG TCA GAG TCT CAG TCA GTG ACA TCA TCA AGA CTG ACT GTG ACT GAC ATG AGT GAC TGA GAC TGT CAT GTC TGA GAG TCG TTT TCG ACT CTC AGA CAT GAC AGT CTC AGT CAC TCA TGT CAG TCA CAG TCA GTC TTG ATG ATG TCA CTG ACT GAG ACT CTG ACT CAC TGA CCC ACT GGC |

**Supplementary Table 2 | Core sequences for the 5' and 3' sets of hairpins.** Colored fonts denote different sections of the hairpin: green = ssDNA overhangs for ligation to the right and left handles, red = XPD poly-dT loading site, blue = hairpin poly-dT tetraloop. Sequences are shown in nomenclature of Integrated DNA Technologies (IDT): Phos = phosphate.

| Oligonucleotide name | Sequence (5'-3') |
|---|---|
| **5' set** | |
| LH base | /5Phos/ CCT GGT TTT TTT TTT AGT CTC AGT CAC TCA TGT CAG TCA CAG TC |
| RH base | /5Phos/ TGA CAT GAG TGA CTG AGA CTC CCA CTG GC |
| Loop | /5Phos/ TCT GAC TCA CTG AGT CAT GTC TGA GAG TCG TTT TCG ACT CTC AGA CAT GAC TCA GTG AGT C |
| Middle complement | /5Phos/ AGA GTC TCA GTC AGT GAC ATC ATC AAG ACT GAC TGT GAC |
| Middle complement off-strand (CA) mismatch | /5Phos/ AGA GTC TCA GTC AGT GAC ATC ATC CAG ACT GAC TGT GAC |
| **3' set** | |
| LH base + middle complement | /5Phos/ CCT GGT TTT TTT TTT TCA GTG AGT CAG AGT CTC AGT CAG TGA CAT CAT CAA GAC TGA CTG TGA C |
| LH base + middle complement off-strand (CA) mismatch | /5Phos/ CCT GGT TTT TTT TTT TCA GTG AGT CAG AGT CTC AGT CAG TGA CAT CAT CCA GAC TGA CTG TGA C |
| RH base | /5Phos/ TCT GAC TCA CTG ACC CAC TGG C |
| Loop | /5Phos/ TGA CAT GAG TGA CTG AGA CTG TCA TGT CTG AGA GTC GTT TTC GAC TCT CAG ACA TGA CAG TCT CAG TCA CTC ATG TCA GTC ACA GTC |
| **Modification inserts** | |
| Middle TT | /5Phos/ AGT CTT GAT GAT GTC ACT GAC TGA GAC |
| Middle CPD | /5Phos/ AGT C{CS-T} GAT GAT GTC ACT GAC TGA GAC |
| Middle abasic | /5Phos/ AGT CT /idSp/ GAT GAT GTC ACT GAC TGA GAC |
| Middle (TC) mismatch | /5Phos/ AGT CTC GAT GAT GTC ACT GAC TGA GAC |
| Middle fluorescein | /5Phos/ AGT CT /iFluorT/ GAT GAT GTC ACT GAC TGA GAC |

**Supplementary Table 3 | Oligonucleotides used to construct hairpins with modifications using piecewise synthesis.** Sequences are shown in nomenclature of Integrated DNA Technologies (IDT), with the exception of the CPD in the 'Middle CPD' oligonucleotide from TriLink Biotechnologies. Refer to Supplementary Fig. 1 for schematics of the oligonucleotides on 5' and 3' sets of hairpins. Abbreviations correspond to the following modifications: Phos = phosphate, {CS-T} = Cis-Syn Thymidine Dimer, idSp = abasic site, and iFluorT = fluorescein.

| **Construct** | **Number of bursts analyzed** | | | | | |
|---|---|---|---|---|---|---|
| | **Unwinding** | | **Rezipping** | | **Rezipping (traversal)** | |
| | **5' set** | **3' set** | **5' set** | **3' set** | **5' set** | **3' set** |
| Unmodified | 81 | 51 | 34 | 28 | 23 | 16 |
| Off-strand mismatch | 39 | | | 23 | | 19 |
| On-strand mismatch | | 51 | 37 | | 25 | |
| CPD | 74 | 48 | 7 | 27 | | 16 |
| Abasic | 108 | 22 | 61 | 11 | 29 | 7 |
| Fluorescein | 39 | 20 | 20 | 13 | 13 | 8 |

**Supplementary Table 4 | Key data statistics.** Number of bursts analyzed under each set of conditions. The unwinding analysis included bursts with processivity of at least 25 bp. The rezipping analysis was restricted to bursts reaching the modification and exhibiting gradual, *bona fide* rezipping (see **Methods**) over at least part of the burst. The rezipping (traversal) analysis used to determine the traversal probabilities included only bursts exhibiting *bona fide* rezipping at 50 bp.
